# Interaction-specific early wheat responses to *Zymoseptoria tritici* share enhanced defense-related transcription during incompatibility

**DOI:** 10.64898/2026.10.07.757330

**Authors:** Marie Unvericht, Frédéric Choulet, Florence Cambon, Florian Rocher, Thierry Langin, Ludovic Bonhomme, Kostya Kanyuka, Cyrille Saintenac

## Abstract

Despite the recent cloning of RLK-encoding resistance genes and the identification of a common point of hyphal arrest at the stomatal level in incompatible interactions, the molecular mechanisms leading to the establishment of wheat resistance against Septoria tritici blotch (STB) remain poorly characterized. Here, we use long-read RNA-sequencing data to compare the early transcriptomic responses of two wheat cultivars carrying major, isolate-specific resistances to *Zymoseptoria tritici* via differential expression, differential transcript usage (DTU), functional- and pathway enrichment analyses. We show that both magnitude and composition of the responses depend strongly on the combination of host cultivar and pathogen isolate and that DTU appears to be uncommon at this infection stage. We do not detect a set of genes exclusively differentially expressed across incompatible interactions, but genes coding for proteins involved in several defense-related processes consistently show higher expression levels in incompatible than compatible interactions, including receptors and ROS homeostatic, antifungal hydrolytic, phenylpropanoid and diterpenoid biosynthetic enzymes. Our results suggest that whilst interaction-specific differences in magnitude and composition may also shape resistance, different major STB resistances may quantitatively enhance overlapping defense machinery, providing a basis for its functional investigation.

## INTRODUCTION

Upon pathogen attack, plant cell-surface receptor-like kinases (RLKs) and receptor-like proteins (RLPs), together with intracellular nucleotide-binding leucine-rich repeat receptors (NLRs), detect pathogen-derived molecules or their effects on host components and activate interconnected signaling networks involving mitogen-activated protein kinases (MAPKs), reactive oxygen species (ROS), Ca^2+^ signals, phytohormones, and multiple transcription factor (TF) families (Couto and Zipfel, 2016; Tsuda and Somssich, 2015). Hence, plant responses to pathogens are associated with extensive transcriptional reprogramming (Chen et al., 2020; Couto and Zipfel, 2016; Tsuda and Somssich, 2015). One post-transcriptional mechanism contributing to responses to pathogens is alternative splicing (AS), which generates multiple transcript isoforms from a single gene (Li et al., 2026; Toh et al., 2025). For example, *Pseudomonas syringae* infection resulted in a more than 100-fold increase of a specific transcript variant of *RESISTANCE TO PSEUDOMONAS SYRINGAE4 (RPS4)* in *Arabidopsis*, whereas a different variant remained stable (Zhang and Gassmann, 2007). These coordinated responses ultimately result in the induction of defense mechanisms such as cell wall reinforcement, production of antimicrobial compounds and, in some interactions, localized cell death (Saur and Hückelhoven, 2021) that restrict the progression of the pathogens.

Septoria tritici blotch, caused by the apoplast-limited fungus *Zymoseptoria tritici*, is an economically important disease of bread wheat (Fones and Gurr, 2015; Savary et al., 2019). *Z. tritici* enters wheat leaves through stomata and subsequently undergoes a prolonged symptomless latent phase characterized by slow growth and low metabolic activity (Brennan et al., 2019; Rudd et al., 2015). After approximately 7-28 days, the fungus transitions to a necrotrophic phase, coinciding with rapid increases in fungal biomass, development of necrotic lesions, and formation of predominantly asexual fruiting bodies, pycnidia, in substomatal cavities (Rudd et al., 2015). Wheat resistance to STB is generally quantitative, although 25 loci with larger effects have been identified and termed *Stb* genes (Brown et al., 2015; Langlands-Perry et al., 2022; Yang et al., 2018; Yang et al., 2026). Only three genes, *Stb6*, *Stb15* and *Stb16q*, have been cloned so far and all encode different subfamilies of RLKs (Hafeez et al., 2025; Saintenac et al., 2018; Saintenac et al., 2021). Resistances conferred by *Stb* genes often depend on the particular combination of host and pathogen genotype (Brown et al., 2015). Furthermore, microscopic studies of wheat cultivars carrying different major STB resistances indicate a degree of mechanistic convergence, with fungal hyphae frequently arrested during or shortly after stomatal penetration (Alassimone et al., 2024; Battache et al., 2022; Battache et al., 2024; Frey et al., 2026; Hansen et al., 2025; Qutb et al., 2024).

Despite the identification of the first STB receptors and the apparent importance of stomatal closure in resistance, the molecular responses following *Z. tritici* perception and leading to the establishment of resistance remain poorly understood. Several transcriptomic studies of wheat infected by *Z. tritici* reported upregulation of genes encoding antioxidant, detoxifying, and hydrolytic enzymes, as well as genes involved in ROS accumulation (Adhikari et al., 2007; Benbow et al., 2020; Gohari et al., 2024; Hansen et al., 2025; Kia et al., 2026). Induction of the phenylpropanoid pathway as well as pathogenesis-related (*PR*) genes was most frequently reported to be associated with resistance (Adhikari et al., 2007; Gohari et al., 2024; Hansen et al., 2025; Kia et al., 2026; Ray et al., 2003; Rudd et al., 2015). However, for some interactions, the opposite trend was also observed or differed based on the investigated timepoint (Benbow et al., 2020; Brennan et al., 2020; Hansen et al., 2025; Rudd et al., 2015). This lack of consensus makes it difficult to identify conserved resistance-related molecular mechanisms. Additionally, analyses were often restricted to single isolate-cultivar combinations or interaction types, with little prior knowledge about the contained host resistance or fungal avirulence genes. A recent study by Alassimone et al. (2024) isolated an Stb6-AvrStb6-specific response using two *Z. tritici* isolates only differing in the presence or absence of the fungal avirulence factor AvrStb6, in combination with direct infiltration of this protein into wheat leaves (Alassimone et al., 2024). This response involved the induction of RLK, cell wall- and stomatal regulation-related genes (Alassimone et al., 2024). Furthermore, the contribution of AS to *Z. tritici*-induced responses of wheat has, to our knowledge, never been investigated. Long-read RNA-sequencing (lrRNA-seq) technologies now allow transcript structures and transcript abundance to be resolved more reliably than previously possible (Wissel et al., 2026). This technological advancement provides an opportunity to explore changes in the relative use of transcript isoforms between conditions via differential transcript usage (DTU) analysis, rather than solely investigating splice-site variation (Lio et al., 2025; Tekath and Dugas, 2021).

Here, we used PacBio Kinnex lrRNA-seq to compare early wheat transcriptomic responses in two cultivars challenged with virulent and avirulent *Z. tritici* isolates. We investigated the extent to which compatible and incompatible interactions share transcriptional responses and whether DTU contributes to their regulation. We identified a shared set of avirulence-associated genes characterized predominantly by higher expression in incompatible than compatible interactions, rather than by genes responding uniquely to infection with avirulent isolates. DTU was rare during early *Z. tritici* infection and mainly involved variation at transcript 5’ ends, often associated with alternative first exon (AFE) usage, rather than AS of internal exons.

## RESULTS

### Bread wheat cultivars Fielder and Jagger possess isolate-specific resistances to *Z. tritici* T48 and IPO90012

We observed strong resistances of the wheat cultivars Fielder and Jagger to *Z. tritici* isolates T48 and IPO90012, respectively, under controlled conditions. Second leaves of Fielder inoculated with T48 and Jagger inoculated with IPO90012 developed barely any disease symptoms, whereas both isolates caused high levels of necrosis and pycnidia on susceptible control L6 (Fig. 1A, B). IPO90012 was highly virulent on Fielder, whereas IPO92006 caused disease on Jagger, demonstrating that both resistances are isolate-specific and, additionally, that the resistance of Jagger to IPO90012 was cultivar-specific. However, disease caused by IPO92006 on Jagger developed more slowly and produced fewer pycnidia than on susceptible L6, suggesting the presence of a low-effect quantitative resistance in Jagger to this isolate (Fig. 1B).

**Figure 1.**
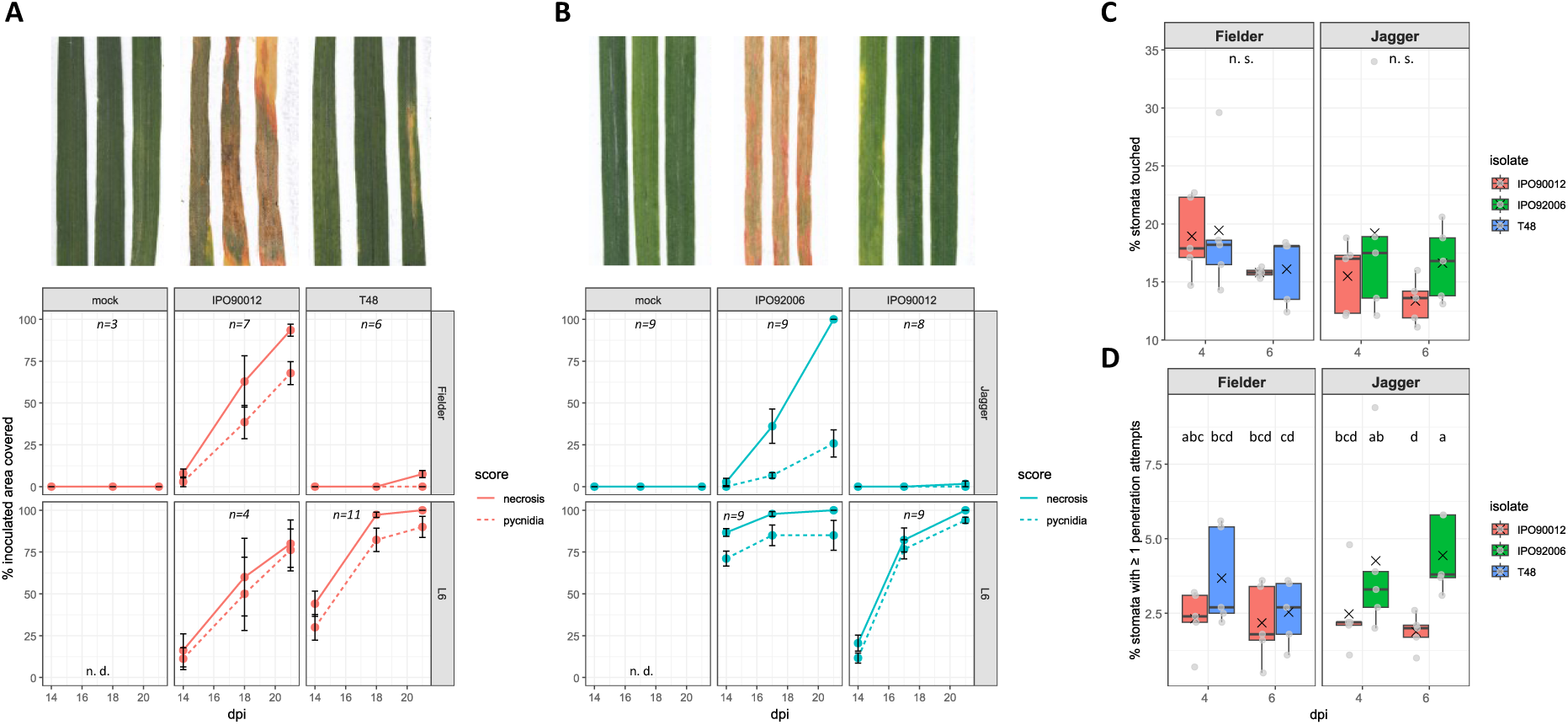
Isolate-specific resistance of *T. aestivum* Fielder to *Z. tritici* T48 and Jagger against IPO90012. (A + B) Representative second Fielder and Jagger leaves at 21 dpi and development of necrosis and pycnidia from 14 to 21 dpi. L6 served as a susceptible control and mock treatment consisted of inoculation buffer (0.05 % Tween-20). Error bars show the standard error. (C) Percentage of stomata contacted by *Z. tritici* hyphae and (D) percentage undergoing at least one penetration attempt under the conditions used for RNA-seq sampling. Crosses indicate means. Different letters indicate significantly different groups according to one-way Van der Waerden test (*χ2* = 14.3, *df* = 7, *p* = 0.046).

To study the molecular responses following initial *Z. tritici* perception, we aimed to identify a timepoint at which stomatal penetration events have just subsided. Therefore, we quantified the proportion of stomata contacted by *Z. tritici* hyphae and undergoing entry attempts 4- and 6-days post infection (dpi; Fig. 1C, D). The proportion of stomata reached by hyphae did not differ significantly between 4 and 6 dpi for any condition, and ranged from 13.4 to 19.4 % across conditions. Stomatal penetration also did not increase significantly between timepoints (Fig. 1D), indicating that most stomatal penetration attempts had already occurred by 4 dpi. However, a small but significant difference in stomatal penetration attempts was observed between IPO92006 and IPO90012 on Jagger at 6 dpi (one-way Van der Waerden test, *χ2* = 14.3, *df* = 7, *p* = 0.046), implying some degree of isolate-specificity.

### Early wheat transcriptomic responses to *Z. tritici* depend on the cultivar-isolate combination

Based on these phenotypic observations, we performed PacBio Kinnex lrRNA-seq using total RNA isolated at 4 dpi from Fielder inoculated with mock, IPO90012^vir^ or T48^avir^, and Jagger inoculated with mock, IPO92006^vir^ or IPO90012^avir^.

Exploration of global gene expression patterns via principal component analysis (PCA) of expressed genes showed that samples separated primarily according to host genotype, reflected by PC1 explaining 56 % of the variation, whereas treatment represented a smaller source of variation along PC2 (approximately 12 %; Fig. 2A). Within Fielder, infection with two isolates produced clearly distinguishable transcriptional profiles, whereas the two infected Jagger conditions clustered more closely. Across contrasts, only 0.3 to 6 % of genes tested after filtering were differentially expressed (25,695 genes tested in Jagger and 28,290 in Fielder), based on consensus differentially expressed genes (DEGs) detected by both edgeR and DESeq2. However, the DEGs generally showed substantial changes in expression, with median edgeR log_2_-fold changes (logFCs) 2.18–2.53 (i. e., fold changes 4.5-5.8; Fig. 2B), and at least 86 % of DEGs were upregulated. Markedly more DEGs and higher logFC extreme values were detected in FLD.T48^avir^ – mock than in the remaining contrasts (Fig. 2B).

**Figure 2.**
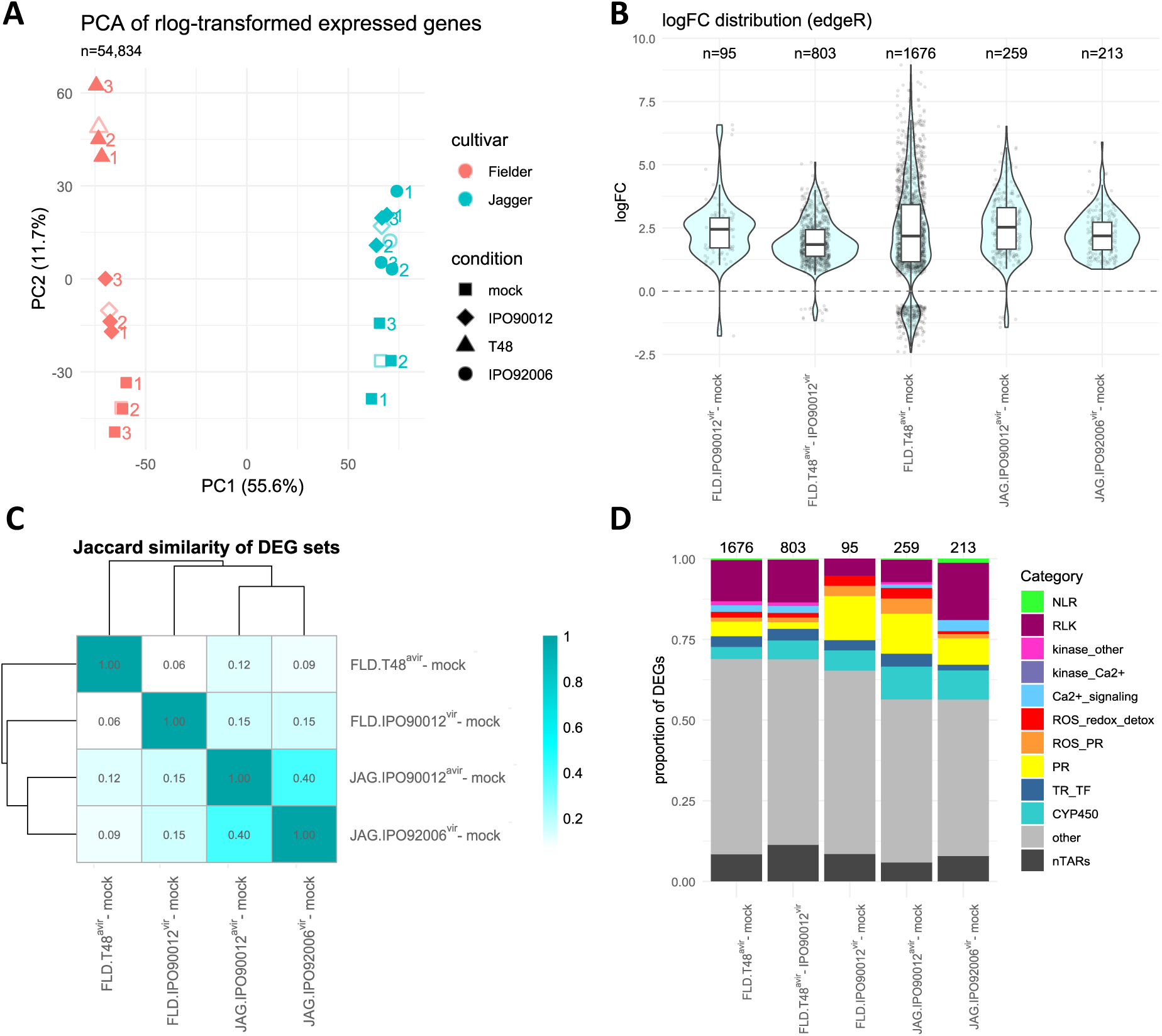
Early wheat transcriptomic responses to *Z. tritici* are dependent on host and pathogen genotypes in both magnitude and composition. (A) Principal component analysis of expressed genes across RNA-seq samples. Both Fielder and Jagger FLNC reads were mapped onto RefSeq_v2.1 for this visualization only. Genes had to be expressed in at least one sample for both cultivars. Unfilled shapes indicate group centroids. (B) Distribution of edgeR log_2_-fold changes across contrasts. FLD = Fielder, JAG = Jagger. (C) Jaccard similarity between DEG sets. (D) Functional composition of DEG sets. NLR, nucleotide-binding leucine rich repeat-containing receptor; RLK, receptor-like kinase; kinase_Ca2+, kinase associated with calcium signaling, ROS_redox_detox, genes associated with ROS scavenging, redox metabolism, detoxification; PR, pathogenesis-related protein; ROS_PR, PR protein associated with ROS metabolism; TR_TF, transcription factor or transcriptional regulator; CYP450, cytochrome P450; nTAR = novel transcriptionally active region. Panels B-D are based on consensus DEGs identified by both DESeq2 and edgeR (FDR/padj ≤ 0.01). JAG.IPO90012^avir^ – JAG.IPO92006^vir^ was omitted in C, D since only a single DEG was identified.

The genes contained in each DEG set also strongly depended on the combination of host cultivar and pathogen isolate. The mean Jaccard similarity (see Experimental Procedures) of DEG sets was only 13.6 %, with the two Jagger responses showing the greatest overlap, corresponding to 40 % of the union of both DEG sets (Fig. 2C). FLD.T48^avir^ – mock showed the lowest similarity to the other contrasts, and only 25 DEGs were shared across all four infection responses (*Supplementary Table S2*). Nevertheless, each DEG set contained genes associated with established plant defense functions, including RLKs, PR proteins, and proteins involved in ROS metabolism (Fig. 2D). NLRs were only rarely differentially regulated. Combination-dependent differences in DEG composition were also evident. For example, both IPO90012 – mock contrasts contained proportionally fewer RLKs and genes associated with Ca^2+^-signaling, but more PR-coding genes, whereas Jagger responses contained a greater proportion of CYP450 class enzymes than other contrasts. Thus, on a global level, both the magnitude and composition of early wheat transcriptional responses to *Z. tritici* depended strongly on host and pathogen genotype.

### Within-cultivar comparisons identify contrasting responses to virulent and avirulent *Z. tritici* isolates

Fielder showed the strongest transcriptional response to avirulent isolate T48 in terms of both DEG number and magnitude of expression changes. 1,059 (63 %) DEGs were exclusive to the FLD.T48^avir^ – mock contrast. The gene whose expression changed the most [edgeR logFC ≈ 8.9] encodes a small, putatively secreted protein with beta-solenoid-like fold, homology to adhesin / collagen-binding domains and overall structural similarity to polygalacturonase inhibiting proteins (*Supplementary Table S2*). iTAK (Zheng et al., 2016) identified 248 RLKs and 63 TFs or transcriptional regulators (TRs) among these DEGs, of which 122 and 30, respectively, were also significantly differentially expressed between T48^avir^- and IPO90012^vir^-infected Fielder. Several defense-associated gene ontology (GO) terms and KEGG pathways were enriched specifically or most strongly in the T48 response, including pathways associated with phytohormones and secondary metabolism (*Supplementary Figures 1 and 2*).

FLD.IPO90012^vir^ – mock contained the fewest (95) DEGs, all of which were also present in FLD.T48^avir^ – mock, and no unique virulence-specific DEGs were detected. Only seven RLKs and three TFs/TRs were identified within this set, and no KEGG pathways and few GO terms were significantly enriched. Processes associated with cell-wall macromolecule catabolism, chitin catabolism and chitinase activity, which were among the most strongly enriched GO terms in all other contrasts (average gene ratios (significant/annotated) of 0.4, 0.5 and 0.5) were not significantly enriched in Fielder infected with IPO90012 (*Supplementary Figure 2*). Two putative lysine motif (LysM) RLKs showed slightly negative, although non-significant, logFCs following IPO90012 infection, but were strongly induced by T48 (Fig. 3A, and *Supplementary Figure 3*). Both showed the highest similarity to Arabidopsis LYK5 (AT2G33580; *e*-values = 3×10^-96^ and 2×10^-96^, 52 % identity, and 45 % coverage), known to bind chitin and trigger immunity (Cao et al., 2014).

**Figure 3.**
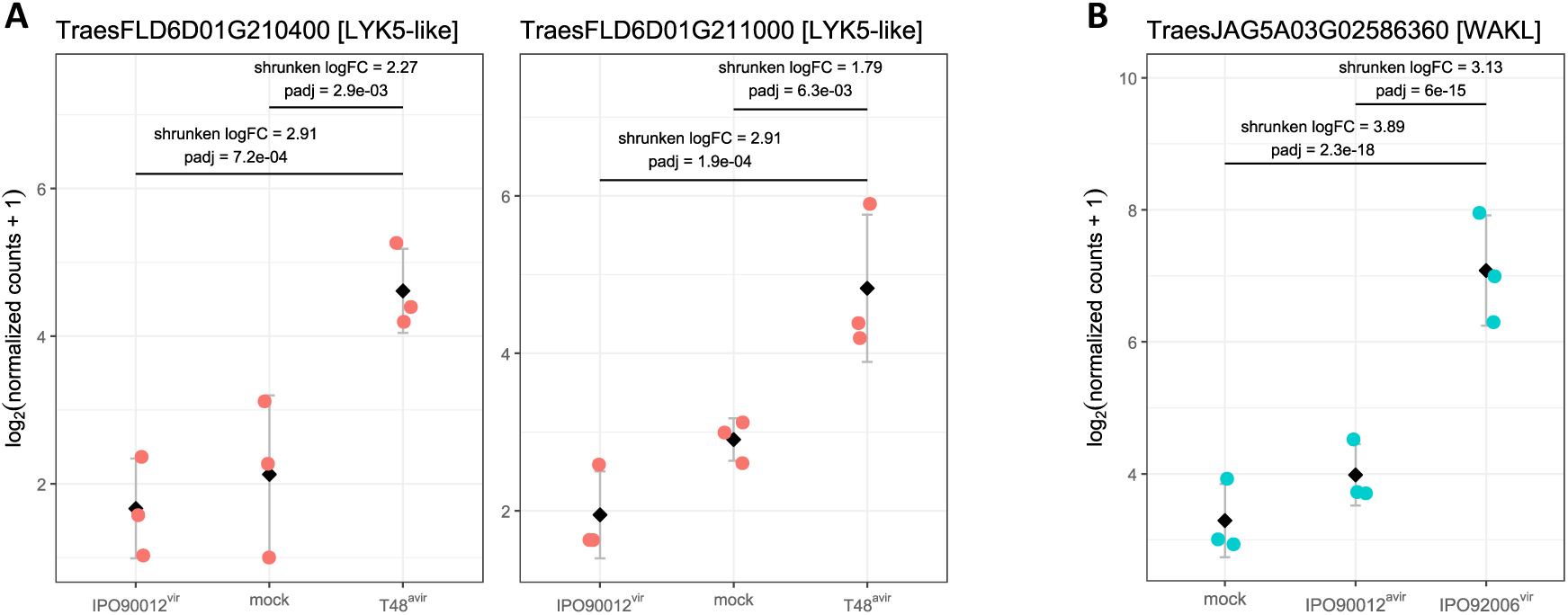
Selected receptor-like kinase genes showing contrasting responses to virulent and avirulent *Z. tritici* isolates. (A) Two putative Fielder LYK5 homologs induced following infection with avirulent T48 but not with the virulent isolate IPO90012. (B) *TraesJAG5A03G02586360*, encoding a WAKL, was the only significant DEG between the incompatible Jagger–IPO90012 and compatible Jagger–IPO92006 interactions, and was strongly induced by IPO92006. Points represent independent biological replicates, diamonds indicate mean expression, and error bars show standard deviation. Expression was normalized using the DESeq2 median-of-ratios method and log₂-transformed for visualization. DESeq2 log2-fold changes were shrunken using the apeglm method.

While a strong difference was observed between Fielder compatible and incompatible interactions, Jagger showed more similar responses to the two isolates (s. Fig. 2B, C). CYP450-coding genes were among the most strongly induced genes in both contrasts, whereas PR proteins (PR-1 and PR-5), a phenylalanine ammonia lyase and an indole-2-monooxygenase-like protein were among genes significantly upregulated in the IPO90012^avir^, but not the IPO92006^vir^ response. The most strongly induced DEG unique to the IPO92006^vir^ response coded for a putative glutamate receptor (*Supplementary Table S2*). iTAK identified 19 and 41 RLKs in JAG.IPO90012^avir^ – mock and JAG.IPO92006^vir^ – mock DEG sets, respectively. Despite these differences, only a single gene, TraesJAG5A03G02586360, was significantly differentially expressed between the two infected conditions (Fig. 3B; *Supplementary Figure 4*). This gene encodes a wall-associated kinase-like protein (WAKL) and was approximately nine-fold more highly expressed in the compatible Jagger–IPO92006 interaction than in the incompatible Jagger–IPO90012 interaction.

### A shared avirulence-associated response is quantitative rather than qualitative

A major objective of this study was to determine whether different incompatible *Z. tritici* interactions share transcriptional responses that are absent from compatible interactions. Several biological processes were enriched in both FLD.T48^avir^ – mock and JAG.IPO90012^avir^ – mock but not in the corresponding virulent-isolate responses (Fig. 4A). These included response to oxidative stress, glutathione metabolism, systemic acquired resistance, diterpenoid biosynthesis and defense response to fungus. Genes whose expression changes underlie these enrichments included peroxidases, catalase, glutathione-S-transferases (GSTs), lipid-transfer proteins, NPR-like proteins, terpene synthases and several classes of PR proteins. *PR-1*, *PR-3*, *PR-4*, *PR-5*, *PR-8* and *PR-9* genes tended to be more numerous and more strongly induced in incompatible interactions, with *PR-2* (β-1,3-glucanases), *PR-3* (chitinases) and *PR-9* (peroxidase) being particularly abundant among DEGs (*Supplementary Figure 5*). Glutathione transferase activity and sequence-specific DNA binding were among molecular functions shared between the incompatible interactions, and several KEGG pathways, including diterpenoid biosynthesis, MAPK signaling, glutathione metabolism, cysteine and methionine metabolism and phenylpropanoid biosynthesis, were enriched in both avirulent-isolate responses (Fig. 4B). Notably, a genome-wide ranked GO analysis identified similar terms as enriched in the same direction in both avirulent-isolate versus virulent-isolate contrasts (*Supplementary Information S1*; *Supplementary Table S6*).

**Figure 4.**
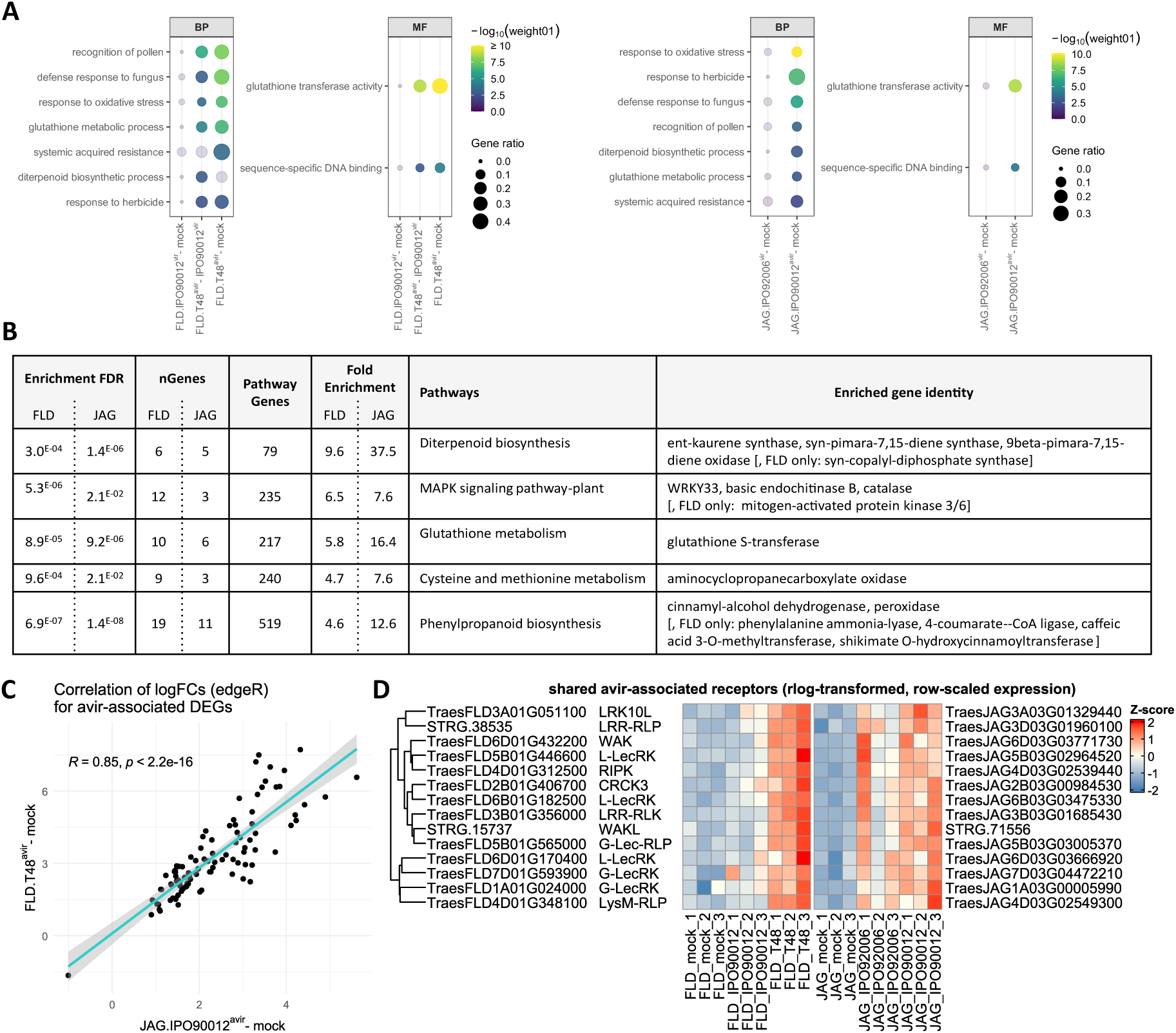
Shared avirulence-associated genes and defense processes across cultivar-isolate combinations. (A) Biological processes (BP) and molecular functions (MF) enriched in the Fielder (left) and Jagger (right) DEG sets. Non-bold dots indicate non-significant terms. Gene ratio represents the number of significant genes divided by the number of annotated genes. (B) KEGG pathways enriched in both incompatible interactions. Enrichment FDR: False discovery rate (FDR)-adjusted *P* value for pathway enrichment, nGenes: Number of significant genes in this pathway in the respective DEG set, Pathway Genes: Total number of genes in the KEGG pathway, Fold Enrichment: Ratio of the proportion of pathway genes in the DEG set to that expected based on the Jagger gene background. (C) Pearson correlation between avirulent-isolate versus mock log_2_-fold changes of the 96 shared avirulence-associated DEGs in Fielder and Jagger. The shaded area indicates the 95 % confidence interval. (D) Expression of fourteen receptor-coding genes among the 96 shared avirulence-associated DEGs. Expression values were regularized-log (rlog) transformed in DESeq2 and row-scaled for visualization. Numbers appended to sample names indicate biological replicates.

Despite this functional overlap, no DEGs from the direct avirulent isolate versus virulent isolate contrasts were shared between Fielder and Jagger, because *TraesJAG5A03G02586360* was the only significant gene in the Jagger isolate comparison. We therefore found no conserved avirulence-specific DEG set in the strict sense. Additionally, the genome-wide concordance of edgeR logFCs between the two avirulent-isolate versus virulent-isolate contrasts was negligible (Pearson’s *r* = 0.054; Spearman’s *ρ* = −0.010), and only 49.5% of homologous gene pairs between Fielder and Jagger showed the same direction of effect in both cultivars (*Supplementary Information S1*). Yet, concordance increased markedly among genes showing the strongest positive direct effects. We next intersected DEGs from the two avirulent-isolate versus mock contrasts and removed genes that were also significantly differentially expressed in either virulent-isolate versus mock contrast. This identified 96 homologous DEGs shared between Fielder and Jagger (*Supplementary Table S4, Supplementary Figure 6*). These genes generally showed strong induction following inoculation with the avirulent isolates and little or modest induction following infection with the corresponding virulent isolates. We therefore refer to them as shared avirulence-associated DEGs (SAADs). Their avirulent-isolate versus mock logFCs were strongly correlated between the Fielder and Jagger datasets (Pearson’s *R* = 0.85, *p* < 2.2×10^-16^; Fig. 4C), consistent with a shared quantitative response. Although SAADs were not significantly differentially expressed across the avirulent-versus virulent-isolate contrasts, their direct edgeR logFC estimates showed a consistent tendency towards higher expression in the incompatible than compatible interactions (Pearson’s *R* = 0.58*; Supplementary Information S1*). In support for a quantitative expression increase across incompatible interactions, when independently ranking genes in each cultivar using a signed edgeR quasi-likelihood statistic for the avirulent-versus virulent-isolate contrast, the overlap between the strongest positive tails substantially exceeded random expectation: The top 1%, 5% and 10% contained 38, 146 and 297 shared genes, corresponding to 19.7-, 3.0- and 1.5-fold enrichment, respectively (*Supplementary Information S1; Supplementary Table S5*).

The 96 SAADs represented multiple layers of plant immunity. Genes potentially involved in perception included nine RLKs and three RLPs from different subfamilies (Fig. 4D). Interestingly, the known RLK-coding resistance genes *Stb6*, *Stb15* and *Stb16q* were only mildly induced following infection, with only *Stb15* significantly differentially expressed in FLD.T48^avir^ – mock, thus they were not SAADs. Signaling-associated SAADs included two calmodulin-like proteins and two receptor-like cytoplasmic kinases (RLCKs) with high similarity to RPM1-INDUCED PROTEIN KINASE (RIPK) and CALMODULIN-BINDING RECEPTOR-LIKE CYTOPLASMIC KINASE 3 (CRCK3), together with four putative 1-aminocyclopropane-1-carboxylate oxidases, three putative 12-oxophytodienoate reductases 1 and a putative salicylic-acid binding protein 2/METHYL ESTERASE 1 homolog. Transcriptional regulation included two WRKY, three HSF, one bHLH, and one C2H2-type zinc finger TF. Defense-associated SAADs included PR-1, PR-2 and PR-5 encoding genes, four chitinases, eight peroxidases and seven GSTs, together with enzymes potentially involved in diterpenoid and phenylpropanoid metabolism. Thus, although incompatible interactions did not share a discrete qualitative transcriptional response, they showed a consistent tendency towards quantitative enhancement of several common defense-associated genes and pathways.

### Few genes show differential transcript usage at 4 days post inoculation

We next wanted to determine whether the preferential expression of alternative gene transcripts could contribute to or differ from the gene-level expression changes observed in wheat during early *Z. tritici* infection. Thus, we *de novo* annotated and quantified transcripts in the Jagger and Fielder datasets to perform DTU analysis using DTUrtle (see Experimental Procedures; *Supplementary Table S1*). The majority of expressed genes were represented by a single isoform, corresponding to 67 % of genes in Fielder and 74 % in Jagger. In Jagger, most transcripts were classified by SQANTI3 as full-splice matches (53 %), followed by intergenic transcripts (19 %) and novel-not-in-catalog transcripts (14 %; for SQANTI transcript terminology, see Tardaguila et al., 2018). In Fielder, the largest categories were novel-not-in-catalog (32 %), intergenic (31 %) and full-splice match transcripts (25 %). The difference between cultivars is likely influenced by the existing Fielder annotation (Sato et al., 2021), which contains only one transcript per gene and in many cases lacks annotated UTRs.

DTU was detected in only a very small proportion of the 8,315 and 13,626 genes tested in Jagger and Fielder after filtering (see Experimental Procedures), respectively (Fig. 5). In Jagger, *STRG.19300* (associated with *TraesJAG3A03G01440520*; for genomic coordinates see *Supplementary Data S3 and S5*) was the only significant DTU gene and was detected in the IPO92006^vir^ – mock contrast. *STRG.19300* encodes a WAKL containing extracellular GUB_WAK and WAK_assoc domains but lacking the EGF repeats characteristic of canonical WAKs. Its four annotated transcripts differ in AFE usage (Fig. 5A), and infection strongly increased expression of the major isoform whereas expression of the other isoforms remained low. A similar, but non-significant, expression pattern was observed following infection with IPO90012^avir^. *TraesJAG3A03G01440520* was not detected as a DEG in the gene-level analysis.

**Figure 5.**
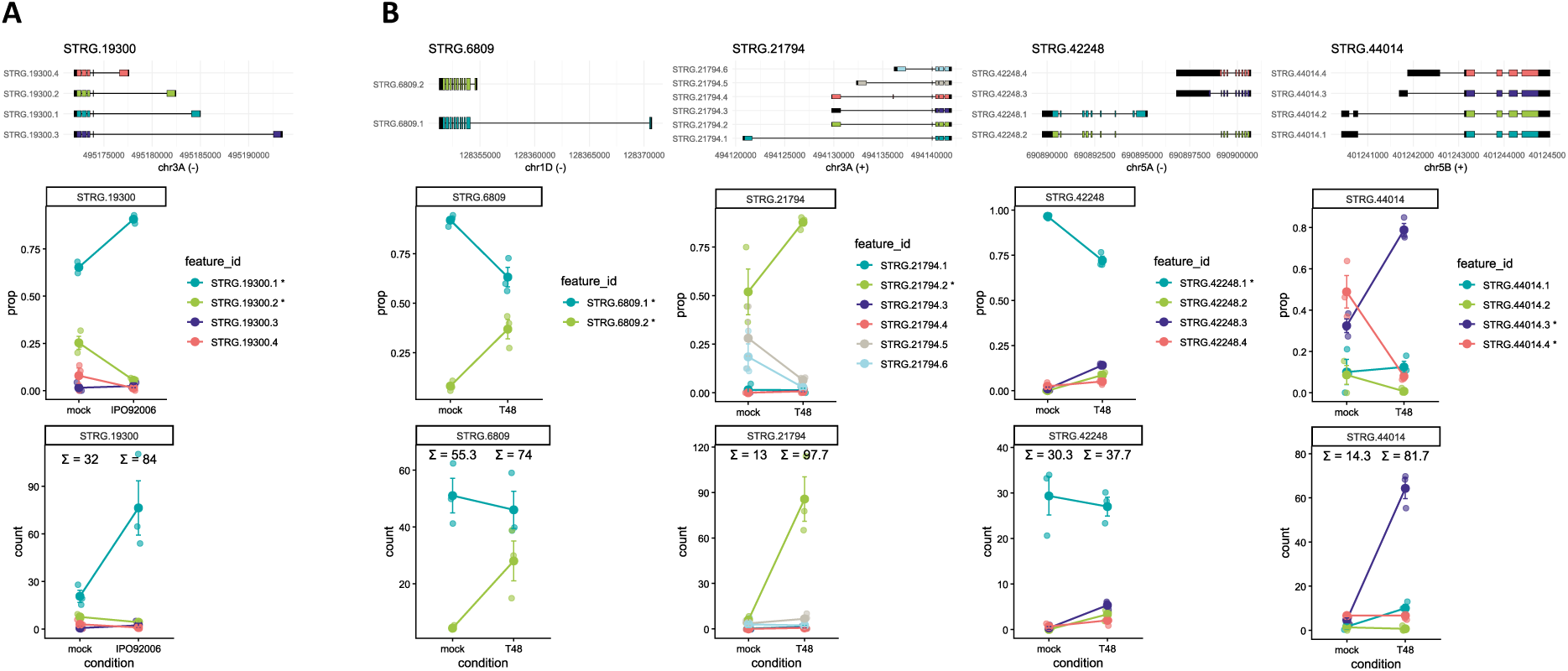
Few genes show significant differential transcript usage 4 dpi after *Z. tritici* infection. Significant DTU genes detected in (A) Jagger for IPO92006^vir^ – mock and (B) Fielder for T48^avir^ – mock. Upper panels show transcript models annotated using StringTie3; colored regions indicate coding sequence and black regions indicate untranslated regions according to SQANTI3. Note that 5’/3’ orientation of transcripts differs according to the strand (+ versus -). Middle panels show the relative contribution of individual transcripts to total gene expression, and lower panels show transcript counts. Values above conditions indicate average total gene counts and asterisks indicate significantly changed transcripts. Transcripts *STRG.19300.3/4*, *STRG.21794.1/3/4* and *STRG.44014.2* were filtered out before statistical analysis but are shown for completeness.

Four genes, *STRG.6809*, *STRG.21794*, *STRG.42248* and *STRG.44014*, displayed significant DTU in Fielder following infection with T48^avir^ (Fig. 5B). Each showed a similar but non-significant trend following IPO90012^vir^ infection. *STRG.6809* (associated with *TraesFLD1D01G138700*) encodes a predicted chloroplastic NADP-dependent malic enzyme. *STRG.6809.1* represents ∼90 % of expression in mock-treated plants, whereas infection shifted expression towards *STRG.6809.2*, which contains a substantially shorter first intron. *STRG.21794* (associated with *TraesFLD3A01G288200*) is most likely homologous to *STRG.19300* in Jagger; despite differences in transcript annotation and orientation on chromosome 3A, their four major alternative proteins share over 99 % sequence identity with Jagger and shared similar transcript trajectories during infection. *STRG.42248* (partially associated with *TraesFLD5A01G561300*) contains a transcript linking genes with similarity to a palmitoyl protein thioesterase and HOMOLOG OF YEAST OXIDASE ASSEMBLY 1 (OXA1). This transcript was supported by 15 FLNC reads across several samples and retains an intact reading frame, making it a candidate fusion transcript that will require independent validation. *STRG.44014* (associated with *TraesFLD5B01G229700*) encodes a putative hypersensitive-induced reaction protein (HIR); all four isoforms share the same coding sequence but differ in their 5’ UTRs, with infection strongly inducing *STRG.44014.3,* which contains one of the shortest 5’ UTR variants. Only *STRG.21794* and *STRG.44014* showed DTU and were also detected as DEGs in Fielder, either for only the FLD.T48^avir^ – mock contrast (*TraesFLD3A01G288200*) or for both infected conditions (*TraesFLD5B01G229700*). Overall, significant DTU was detected in only five genes at 4 dpi and predominantly involved variation at transcript 5’ ends, often associated with AFEs, rather than alternative splicing of internal exons. Additionally, some genes displaying DTU under infection were not DEGs, highlighting that changes in transcript usage can occur without substantial changes in overall gene expression.

## DISCUSSION

Here, we compared early transcriptomic responses of two bread wheat cultivars to virulent and avirulent *Z. tritici* isolates using lrRNA-seq. Using the PacBio Kinnex data, we determined that approximately 59 K annotated genes were expressed per condition, corresponding to roughly 41 % of Jagger and 51 % of Fielder genes (*Supplementary Table S1*). We were also able to detect a substantial number of nTARs (Fielder: ∼118 K annotated genes, ∼26 K nTARs; Jagger: ∼140 K annotated genes, ∼12.5 K nTARs; *Supplementary Table S1*), demonstrating the potential of improving existing gene annotations using long-read RNA-seq. Notably, many of these nTARs were differentially expressed (Figure 2D), including many RLKs (*Supplementary Figures 4 and 5*). Three main conclusions emerge from the study. First, early transcriptional responses were strongly dependent on the particular host cultivar–pathogen isolate combination. Second, we found no conserved set of genes specifically differentially expressed during incompatible interactions and little genome-wide concordance in direct avirulent-isolate versus virulent-isolate expression effects (*Supplementary Information S1*). Nevertheless, a consistent pattern was evident among genes showing the strongest increases in expression, and a set of 96 avirulence-associated DEGs as well as several defense processes showed a similar tendency towards stronger induction in incompatible than compatible interactions. Third, at 4 dpi, significant DTU was detected in only five genes, with most detected events involving variation at transcript 5′ ends rather than alternative splicing of internal exons. Together, these results suggest that different STB resistances may share specific defense processes while retaining substantial quantitative and genotype-specific differences in the underlying transcriptional response.

The transcriptional responses observed in individual compatible interactions further illustrate the strong dependence of these responses on host and pathogen genotype. Fielder showed an unusually weak transcriptional response to virulent IPO90012, with few induced RLKs and little enrichment of chitin-associated defense processes. Two *LYK5*-like genes were strongly induced by avirulent T48 but not by IPO90012. Wheat TaCEBiP and TaLYK5 proteins contribute to chitin-triggered immunity (Liu et al., 2023), whereas *Z. tritici* LysM effectors can interfere with chitin recognition (Lee et al., 2014; Marshall et al., 2011). It is therefore possible that reduced activation of chitin-associated signaling contributes to the compatible Fielder– IPO90012 interaction, although the non-significant expression changes in the two *LYK5*-like genes do not by themselves establish active suppression by the pathogen. Similarly, the strong induction of a *WAKL* during the compatible Jagger–IPO92006 interaction could reflect either pathogen-induced host response modulation or host responses associated with a partial resistance of Jagger to this isolate. Functional analyses will be required to distinguish these possibilities.

### Potential mechanisms contributing to hyphal arrest in the substomatal cavity

Previous studies suggest that wheat plants carrying different major STB resistance genes frequently block *Z. tritici* penetration at the stomata (Alassimone et al., 2024; Battache et al., 2022; Battache et al., 2024; Frey et al., 2026; Hansen et al., 2025; Qutb et al., 2024). For incompatible interactions involving *Stb6*, *Stb9*, or *Stb16q*, *Z. tritici* challenge transiently increased stomatal closure, possibly limiting fungal penetration (Battache et al., 2022; Frey et al., 2026; Ghiasi Noei et al., 2022). Supporting this observation, Alassimone et al. (2024) reported a transcriptional induction of several stomatal closure-related genes during early *Stb6*-mediated resistance, which we could not detect for the resistances studied here (discussed below). Battache et al. (2024) found, that where stomatal penetration occurred, fungal growth was subsequently arrested within or near the substomatal cavity, and direct infiltration of spores into leaves of resistant cultivars largely restricted disease to the infiltrated region. These observations support the existence of a second defense layer within the apoplast (Battache et al., 2024). Our results identify several processes that could contribute to such a hostile apoplastic environment.

Phenylpropanoid biosynthesis was strongly associated with incompatible interactions, including induction of cinnamyl-alcohol dehydrogenases (CAD) and peroxidases (Fig. 4B), consistent with increased lignin production. Cell wall reinforcement through lignification could physically restrict fungal growth, reduce nutrient availability, and cause phenolic toxicity during incomplete lignin polymerization (Tiwari et al., 2025; Yu et al., 2026). Higher *Z. tritici*-induced lignin accumulation has previously been reported in resistant Chinese Spring compared with susceptible Obelisk (Seybold et al., 2020), whereas three *CAD* genes were downregulated during early infection of susceptible Riband (Rudd et al., 2015). Our results also implicate regulation of ROS homeostasis, through avirulence-associated GSTs, catalases and peroxidases together with a RIPK homolog potentially linked to RBOHD-dependent ROS production (Li et al., 2021). This is consistent with previous observations that the timing and magnitude of H_2_O_2_ accumulation differ between resistant and susceptible wheat responses to *Z. tritici* (Ghiasi Noei et al., 2022; Gohari et al., 2024).

Antifungal hydrolytic enzymes may provide another component of this apoplastic defense. PR-3 chitinases were more strongly represented in incompatible interactions, and several SAADs encoded chitinases, consistent with previous reports of stronger induction of chitinases in resistant cultivars (Adhikari et al., 2007; Gohari et al., 2024). In addition, a chromosome 2D-localized group of SAADs encoded enzymes associated with diterpenoid metabolism and corresponded to a cluster recently linked to the production of wheat diterpenoid aspisosides (Misra et al., 2026). The genes included *TaKSL-D1*, *CYP99A8_2D*, *CYP99A10_2D*, and *CYP99A7_2D*. Misra et al. (2026) found that *TaKSL-D1* is induced by several fungal pathogens, including *Z. tritici*, and that aspisosides have direct antifungal activity against *Fusarium culmorum*, *Magnaporthe oryzae*, *Rhizoctonia solani* and *Botrytis cinerea* (Misra et al., 2026), suggesting that these compounds may also contribute to STB resistance.

### Why are wheat transcriptomic responses to STB so dependent on the cultivar-isolate combination?

Host genotype explained the largest component of variation in our expression data, consistent with previous transcriptomic studies of the wheat–*Z. tritici* interaction (Benbow et al., 2020; Gohari et al., 2024). Previous studies have also reached apparently contrasting conclusions, with resistance associated either with stronger transcriptional activation (Adhikari et al., 2007; Gohari et al., 2024; Ray et al., 2003) or with a more restricted response (Benbow et al., 2020; Brennan et al., 2020). Our results similarly show limited overlap in individual resistance-associated genes between different host–isolate combinations, despite evidence for functional overlap in several defense processes. For example, none of the 247 AvrStb6-specific DEGs reported by Alassimone et al. (2024) corresponded directly to our SAAD set. Nevertheless, ten SAADs showed a similar expression pattern in that study, including a WAKL, a G-type LecRK, a G-type LecRP and the CRCK3 homolog. These genes therefore represent useful candidates for testing whether parts of this response are shared more widely between genetically distinct STB resistances. The limited similarity in transcriptomic responses to *Z. tritici* infection is somewhat surprising, given the shared point of fungal arrest observed for multiple major STB resistances (Alassimone et al., 2024; Battache et al., 2022; Battache et al., 2024; Frey et al., 2026; Hansen et al., 2025; Qutb et al., 2024).

Several factors could contribute to this apparent combination-dependence. First, a common resistance-associated response may be highly localized around stomata and therefore strongly diluted in bulk-leaf RNAseq data. Single-cell or spatial transcriptomic approaches focused on guard cells, subsidiary cells and neighboring tissues may resolve responses that are not detectable at whole-leaf level. Second, successful resistance may depend more strongly on the timing of activation than on the final composition of the response. Temporal differences have previously been implicated in STB resistance (Gohari et al., 2024; Hansen et al., 2025; Ray et al., 2003; Rudd et al., 2015), but cannot be resolved from the single 4 dpi time point analyzed here.

Finally, resistance mechanisms may not be primarily regulated through changes in transcript abundance. Consistent with this, Hansen et al. (2025), who investigated STB resistance in the wheat wild relative *Aegilops cylindrica*, concluded that “there is not a specific immune response that hinders the penetration of the pathogen visible in the transcriptomic response”. Immune signaling is extensively regulated at translational and post-translational levels, including phosphorylation, S-nitrosylation, and ubiquitination (Couto and Zipfel, 2016; Frederickson Matika and Loake, 2014; Gough and Sadanandom, 2021). We detected no DEGs directly associated with canonical stomatal closure pathways, apart from the avirulence-associated RIPK homolog linked to RBOHD-mediated ROS production (Li et al., 2021). Stomatal closure relies strongly on rapid post-transcriptional regulation of preexisting proteins (Balmant et al., 2016; Geiger et al., 2009), making it unlikely that all relevant signaling events would be visible through transcript abundance alone. Phosphoproteomic analysis of early time points following *Z. tritici* perception may therefore provide an important complement to transcriptomics and help resolve the signaling mechanisms that determine successful resistance.

### Limited DTU at 4 dpi and preferential involvement of transcript 5’ ends

The majority of our *de novo* transcript annotations were represented by only a single expressed isoform. This is broadly consistent with previous observations in wheat and suggests that the limited number of detected isoforms is not solely a consequence of incomplete transcript annotation (Yu et al., 2020). However, a recent study detected a substantially higher proportion of multiple-isoform genes in wheat leaves, which could be due to different degrees of stringency when defining isoforms (Dai et al., 2026). We could only detect five genes with significant DTU under *Z. tritici* infection. Because our analysis was restricted to a single time point (4 dpi), however, we cannot exclude more extensive or transient changes in transcript usage at other stages of infection. Zhang et al. reported that only approximately 10–11 % of expressed wheat genes underwent alternative splicing during infection with *Blumeria graminis* f. sp. *tritici* or *Puccinia striiformis* f. sp. *tritici* (Zhang et al., 2019). While alternative splicing and DTU are not directly equivalent, the relatively limited extent of infection-associated alternative splicing reported previously, together with the low frequency of DTU observed here, suggests that, at least at the timepoint examined, infection-induced regulation in wheat may occur predominantly at the gene rather than transcript level. Consistent with this, many differentially expressed multi-isoform genes in our data showed proportional changes across their isoforms. The extensive expansion of gene families in wheat may provide one possible explanation, as a negative relationship between gene-family expansion and alternative splicing has been reported across several organisms and forms the basis of the proposed “functional sharing” model (Yu et al., 2020). Wheat contains three homoeologous genomes together with large families of duplicated defense-related genes, including WAK/WAKL-encoding genes, potentially providing functional diversity at the gene family level without requiring extensive DTU. This remains speculative, but is consistent with the low frequency of DTU observed here.

Notably, most significant DTU events detected in this study involved differences at transcript 5′ ends or alternative first exons rather than alternative processing of internal exons. AFEs frequently arise through alternative promoter use (Chen et al., 2007). The resulting transcript abundances can therefore be partly independent of one another, differing conceptually from classical alternative splicing-driven DTU. AFE events are relatively rare, affecting approximately 5 % of *Arabidopsis* or rice, and 1 % of wheat genes (Chen et al., 2007; Yu et al., 2020). Thus, their predominance among the DTU events detected here is notable. Interestingly, Zhang et al. (2019) found that *Bgt*-and *Pst*-associated alternative splicing events in wheat frequently affected transcript extremities rather than internal exons. Together, these findings suggest that regulation of transcript initiation and 5′ transcript architecture may play a greater role in wheat pathogen-responsive transcript diversity than their overall frequency would suggest.

## EXPERIMENTAL PROCEDURES

### Plant and fungal growth conditions

Four grains per 10 x 10 cm pot of the bread wheat cvs Fielder, Jagger and a near-isogenic line of Chinese Spring without *Stb6* (L6, susceptible infection control) sown in HUMUSTAR seedling soil (formula N° SEMHUMU, HUMUSTAR TERREAUX, France) were vernalized in a MTR30 growth chamber (Conviron) for 4 days at 6°C and 80 % relative humidity, followed by 10 days under LED lighting (VEGELED Horticulture Profiles, Eos Series) with a 16-h photoperiod at 21/18 °C (day/night). Plants were watered twice a week until inoculation, then fertigated three times a week with approx. 170 mL nutrient solution per pot. The solution contained 10 kg FERTIPLANT 10 – 10 – 30 + 5 MgO (PLANTIN, France), 14 kg calcium nitrate (YaraLiva^TM^ CALCINIT^TM^, Yara Norge AS) and 2 L 52,5 % nitric acid (VWR International SAS, France) per 100 L total volume. *Zymoseptoria tritici* T48 (IN95-Lafayette-1196-WW 1-4), IPO90012 and IPO92006 were cultured for 4 days in YG medium containing 25 µg/mL each ampicillin and streptomycin sulfate at 20°C and 180 rpm, followed by 3 days on YPD agar with the same antibiotics at 20°C.

### Plant inoculation and disease assessment

*Z. tritici* cultures were suspended in 0.05 % (v/v) Tween-20 and concentrations adjusted to 1 × 10^6^ spores / ml for inoculation. Pure 0.05 % (v/v) Tween-20 served as the mock treatment. A circa 6 cm long section on the second leaves of the 14-day-old wheat seedlings (growth stage Z12 – Z13) was marked and paintbrush-inoculated with two series of six brush strokes, drying between series. Inoculated plants were well watered and covered with transparent plastic bags for 3 days to facilitate infection, then cultivated as described above. Plants not sampled for RNA extraction were kept to assess disease at 14, 17 or 18 and 21 dpi by visual estimation of the percentage of inoculated leaf area covered with necrosis and pycnidia, respectively.

### Sample preparation

At 4 dpi, after 10 h illumination, inoculated leaf sections were harvested in liquid nitrogen. For each condition (cultivar-isolate combination), one biological replicate comprised pooled leaf tissue from nine plants. The complete growth and inoculation experiment was performed independently three times, yielding three biological replicates per condition. RNA was extracted using the QIAGEN RNeasy Plus Mini Kit following the manufacturer’s instructions, RNA concentration was measured using an Invitrogen Qubit 4 fluorometer (Thermo Fisher Scientific) and RNA integrity was assessed by gel electrophoresis.

### RNA sequencing and read mapping

Library preparation using the Kinnex full-length RNA kit (Pacific Biosciences of California, Inc. (PacBio), 2024a) and cDNA sequencing on one Revio SMRT cell per cultivar (corresponding to nine samples) was performed by the INRAE platform GENTYANE (https://gentyane.clermont.inrae.fr). PacBio Kinnex builds on the MAS-ISO-seq technology described by Al’Khafaji et al. (2024). HiFi reads had a mean length of 12.96 kb or 13.96 kb and an N50 of 13.03 kb or 14.07 kb for Fielder and Jagger datasets, respectively, with a median read quality of Q36 or Q34. Following preprocessing and deconcatenation, ∼6-7 million FLNC reads per sample were obtained (*Supplementary Table S1*). FLNC were mapped to the published Jagger (PGSBv2.1, INSDC Assembly GCA_903993795.1) (Walkowiak et al., 2020) and Fielder (201216_Fielder_pseudomolecules_V1+unanchored_contigs.fasta) (Sato et al., 2021) genomes using pbmm2 (Pacific Biosciences of California, Inc. (PacBio), 2024b) (version 1.16.0) with PacBio ISOSEQ preset and --min-id-perc 99. 95–98 % of reads mapped to the respective wheat genome, and under 0.1 % to *Z. tritici* IPO323 (MYCGR v2.0, NCBI Assembly GCF_000219625.1; *Supplementary Table S1*) (Goodwin et al., 2011). For the PCA in Figure 4 only, FLNC reads of both Fielder and Jagger datasets were mapped onto IWGSC CS RefSeq_v2.1 (GCA_018294505; *Supplementary Data S7*) (Zhu et al., 2021).

### Differential gene expression analyses

#### 1. Annotation of nTARs

Novel transcriptionally active regions (nTARs) were defined as transcribed regions that do not overlap previously annotated genomic features (Zeller et al., 2009; Zhu et al., 2014). Within each cultivar, BAM files from all samples were merged prior to running StringTie (version 3.0.3) (Shinder et al., 2026a; Shinder et al., 2026b) in reference-guided (option -G) long-read mode (option -L). Transcripts not overlapping any annotated genes were then identified using bedtools intersect ( v2.31.1; option -v) (Quinlan and Hall, 2010) and classified as nTARs. nTARs were concatenated to the reference annotation and reads aligned to each annotated gene or nTAR by pbmm2 were counted as one count per assigned read. Raw gene-wise counts tables for DEG analyses are provided as *Supplementary Data S1 and S2*.

#### 2. DEG testing with edgeR and DESeq2

DEG analysis in R (version 4.4.3) (R Core Team, 2025) was performed separately for each cultivar on annotated genes and nTARs, using gene-level counts generated in the previous step with both edgeR (version 4.4.2) (Chen et al., 2025) and DESeq2 (version 1.52.0) (Love et al., 2014). Raw counts were filtered using edgeR’s filterByExpr() with min.count=15; the resulting matrix was used as input for both methods to ensure a common gene universe. Additional filtering was therefore omitted in edgeR, and independentFiltering=FALSE was set in DESeq2’s results(). Unless otherwise specified, default parameters were used. The design formula for both tools was ∼repetition + treatment, whereby treatment corresponded to mock, avirulent or virulent isolate inoculation. Normalization and dispersion estimation were performed with calcNormFactors(), followed by estimateDisp() for edgeR, and DESeq() for DESeq2. Differential expression was tested using edgeR’s quasi-likelihood negative binomial GLM framework, with model fitting using glmQLFit() followed by hypothesis testing using glmQLFTest(), and DESeq2’s Wald test with results(). The tested pairwise contrasts corresponded to virulent isolate vs. mock, avirulent isolate vs. mock and avirulent isolate vs. virulent isolate. Genes with FDR ≤ 0.01 (edgeR) / padj ≤ 0.01 (DESeq2) were considered significantly differentially expressed. DESeq2 log_2_FoldChanges were shrunken using the apeglm method (Zhu et al., 2019) prior to visualization and ranking, but after significance testing.

#### 3. Definition of consensus DEGs and visualization

For subsequent analyses, only contrast-wise consensus DEGs, significant for a given contrast in both edgeR and DESeq2 analyses, were considered. Per-contrast consensus DEGs can be found in *Supplementary Table S2*. Results were visualized using ggplot2 (version 4.0.3) (Wickham, 2016). Heatmaps were generated with ComplexHeatmap (version 2.22.0) from edgeR logFCs or row-scaled DESeq2 rlog-transformed counts, using Euclidean distance and complete-linkage clustering. The similarity of two DEG sets A and B was quantified using the Jaccard index (Jaccard, 1901), calculated as *J*(*A*, *B*) = |*A* ∩ *B*|⁄|*A* ∪ *B*|. Jaccard distances (1 – Jaccard index) were then hierarchically clustered using average linkage.

#### 4. Classification of DEGs

Categories for annotated DEGs, except kinases and TR_TFs, were defined by the presence of characteristic PFAM domains (InterProScan version 5.76-107.0; Jones et al. (2014)) in their protein sequences, as listed in *Supplementary Table S3*. Putative PR proteins were BLASTed against a custom UniProt-based database of known PR proteins to confirm their identity (*Supplementary Table S3*). Kinases, TFs and TRs were detected using iTAK (Zheng et al., 2016) either only in annotated genes (Fig. 2D) or also among nTARs, for numbers mentioned in the main text. For nTARs, iTAK was provided with genomic sequences, as we did not predict coding phases for nTARs. Classification results are provided in *Supplementary Table S3*.

#### 5. Functional enrichment analyses of DEG sets

GO terms for all genes subjected to DEG testing were recovered from ensembl through biomaRt (version 2.62.1) (Durinck et al., 2009), directly using the tajagger_eg_gene dataset for Jagger and using tarefseqv2_eg_gene for Fielder after recovering RefSeq_v2.1 homologs. Fielder–Jagger/RefSeq_v2.1 homologs were identified by chromosome-wise reciprocal best BLAST hits on CDS sequences using ≥80% coverage and identity. Enrichment analyses of DEG sets against all genes subjected to DEG testing were performed ontology-wise using topGO (version 2.58.0) (Alexa et al., 2006) with nodeSize=10. Significance testing used the weight01 algorithm with fisher statistic, and significance was defined as weight01 ≤ 0.01. For KEGG enrichment, Fielder DEG identifiers were converted to Jagger homologs using the reciprocal-BLAST mapping, and DEG sets were enriched with ShinyGO (version 0.85.1) (Ge et al., 2020) using the Jagger gene set as background and default parameters.

#### 6. Definition of SAADs

Shared avirulence-associated DEGs were defined as the intersection of significant DEGs for the Fielder and Jagger incompatible interactions, subtracted by those genes also significant for either compatible interaction. DEGs were translated into the Jagger namespace using the reciprocal-BLAST mapping (see above) beforehand.

### Differential transcript usage analysis

#### 1. *De novo* transcript annotation and quantification

Within-cultivar *de novo* transcript annotation was chosen to ensure a comparable transcript definition between cultivars. We deployed a pipeline of StringTie (version 3.0.3) to annotate transcripts, SQANTI3 (5.5.4) for quality control and filtering, and IsoQuant (3.13.0) to obtain counts per transcript based on the filtered transcriptome. StringTie was run reference-free (unspecified -G) because reference-guided annotation resulted in substantial transcript loss during IsoQuant processing. BAM files of all samples within one cultivar were merged prior to running StringTie with -L and --fr. SQANTI3 was provided with the respective Fielder or Jagger reference genome FASTA and annotation files (see read mapping). Transcript models were evaluated using its default rules-based filtering criteria, and SQANTI3 rescue was performed in automatic mode with requantification enabled. SQANTI3’s isoforms_rescued.gtf and the respective reference genome FASTA file were provided to IsoQuant, together with per-sample BAM files, to obtain transcript-level counts using --data_type pacbio_ccs. For SQANTI classification terminology, see (Tardaguila et al., 2018). nTAR and *de novo* transcript identifiers use the same StringTie naming convention but were generated independently; identical identifiers therefore do not denote the same features.

#### 2. DTU testing with DTUrtle

DTU analysis was performed separately per cultivar. The transcript-level quantification data described in the previous section was used as an input for DTUrtle together with SQANTI3’s isoforms_rescued.gtf annotation (*Supplementary Data S3 – S6*). DTUrtle uses DRIMSeq for DTU analysis and stageR for multiple-testing correction (Tekath and Dugas, 2021). To avoid read-length based normalization, count matrices were read in using read.delim() instead of DTUrtle’s import_counts(). To retain a similar gene universe between contrasts, a custom sequential filtering strategy was applied: Transcripts were first required to be expressed in ≥2 samples and to contribute ≥5% of gene expression in ≥3 samples, after which genes were retained only if ≥2 transcripts remained. Subsequently, in run_drimseq(), filtering_strategy was specified as “own” and all min_[…] parameters were set to zero. The tested contrasts were the same as described for the DEG analysis, with each contrast analyzed separately using run_drimseq(). posthoc_and_stager() was performed with default parameters (ofdr=0.05, posthoc=0.1) to identify genes showing an overall significant change in transcript proportions and, where possible, specific transcripts significantly driving this change. Plots were generated using ggtranscript (version 1.0.0) (Gustavsson et al., 2022) and ggplot2.

### Quantification of stomata reached and subjected to penetration

For microscopic analysis, plants were grown and inoculated under the same conditions as described above. At 4 and 6 dpi, approximately 3-cm-long leaf sections were collected from 5 leaves per cultivar-isolate combination, stained with 0.1% Calcofluor White M2R (Sigma-Aldrich) and observed with a Leica DM5500 B fluorescent microscope as described previously (Battache et al., 2022; Battache et al., 2024; Frey et al., 2026). For each leaf, five random Z-stacks comprising 25-35 images were acquired, whereby one stack captured a visual field of on average 37 stomata. Z-stacks were analyzed in Fiji (ImageJ 1.54p; Java 21.0.7) (Schindelin et al., 2012). For each stack, proportions of stomata contacted by *Z. tritici* and undergoing ≥ 1 penetration attempt were calculated relative to total stomata. The average proportion value calculated from the 5 Z-stacks per leaf was then considered as one biological replicate for plotting and statistical testing. Stomatal penetration attempts were identified by hyphae that appeared to extend partially into stomatal pores, often with appressorium-like hyphal thickenings on the ostioles (Battache et al., 2022; Frey et al., 2026). *Supplementary Figure 7* shows examples of the microscopic observations. One-way non-parametric Van der Waerden tests (Conover, 1999) were performed across all isolate-cultivar-dpi combinations using the waerden.test() function from the agricolae (version 1.3-7) package (de Mendiburu Delgado, 2023).

## ACKNOWLEDGEMENTS

We would like to thank INRAE VégéPôle, especially Richard Blanc, for helping with the growth chamber assays, and Vincent Pailler, Elodie Belmonte and Véronique Gautier from INRAE GENTYANE for PacBio Kinnex library preparation, sequencing and pre-processing. We thank Nuren Tasneem for determining structural similarity of TraesFLD5B01G573600 with polygalacturonase inhibiting proteins. *Z. tritici* isolates were kindly provided by Gert Kema and Thierry Marcel. Jagger and Fielder seeds originate from the SeedStor of the John Innes Centre and the National BioResource Project (Japan), respectively. The authors acknowledge the support received from the International Research Center on Sustainable AgroEcosystems (ISITE CAP20-25).

## CRediT AUTHOR CONTRIBUTIONS

• M. U.: Conceptualization, Formal analysis, Investigation, Methodology, Visualization, Writing – original draft

• F. Ch.: Formal analysis, Methodology, Writing – review & editing

• F. Ca.: Investigation

• F. R.: Methodology

• T. L.: Funding acquisition, Supervision, Writing – review & editing

• L. B.: Methodology, Supervision, Writing – review & editing

• K. K.: Conceptualization, Formal Analysis, Supervision, Writing – review & editing

• C. S.: Conceptualization, Funding acquisition, Methodology, Supervision, Writing – review & editing

## DATA AVAILABILITY STATEMENT

Raw sequence (FLNC) data generated in this study has been deposited in the European Nucleotide Archive (ENA) under study accession no. ERP206591, with run accession no. ERR17961755, ERR17962102, ERR17962251, ERR17962268, ERR17962326, ERR17962526, ERR17963012, ERR17963037, ERR17963058, ERR17963133-ERR17963135, ERR17963139 and ERR17963147-ERR17963151. Analysis scripts are available from the authors upon reasonable request.

## Notes

### Competing Interest Statement

The authors have declared no competing interest.

