## Supplementary_Figures_S1-S7 for "Interaction-specific early wheat responses to *Zymoseptoria tritici* share enhanced defense-related transcription during incompatibility"

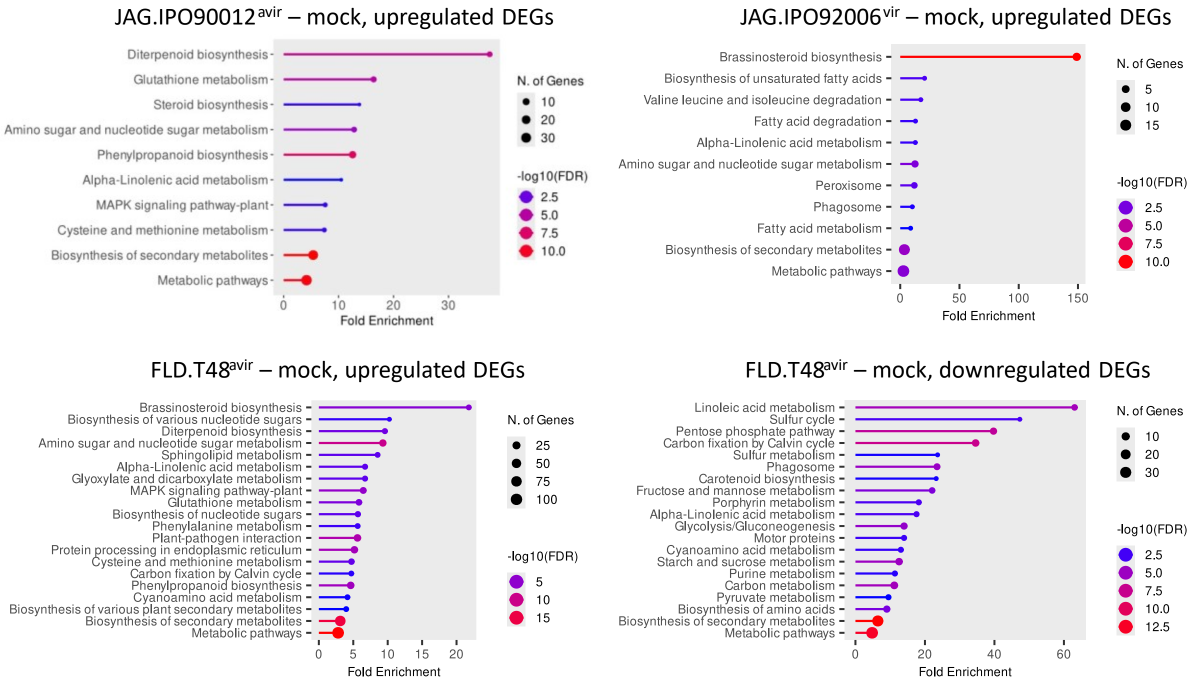


**Supplementary Figure 1: Per-DEG set significantly enriched KEGG pathways.** Enrichment analyses were performed and plots generated using ShinyGO 0.85.1 (Ge et al., 2020; species: tajagger_eg_gene; default parameters).

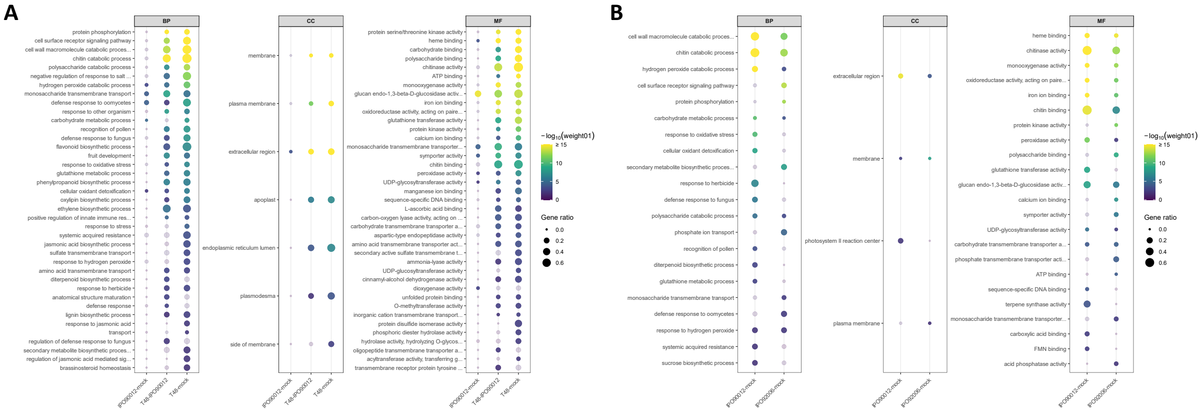


**Supplementary Figure 2: Per-DEG set significantly enriched GO terms for the Fielder (A) and Jagger (B) data.** BP = biological process, CC = cellular component, MF = molecular function. Non-bold dots indicate non-significance. Gene ratio = Significant/Annotated. GO terms were recovered from ensembl using R/BiomaRT (tajagger_eg_gene dataset) and enriched using R/topGO (weight01 algorithm [weight01 ≤ 0.01], fisher statistic).

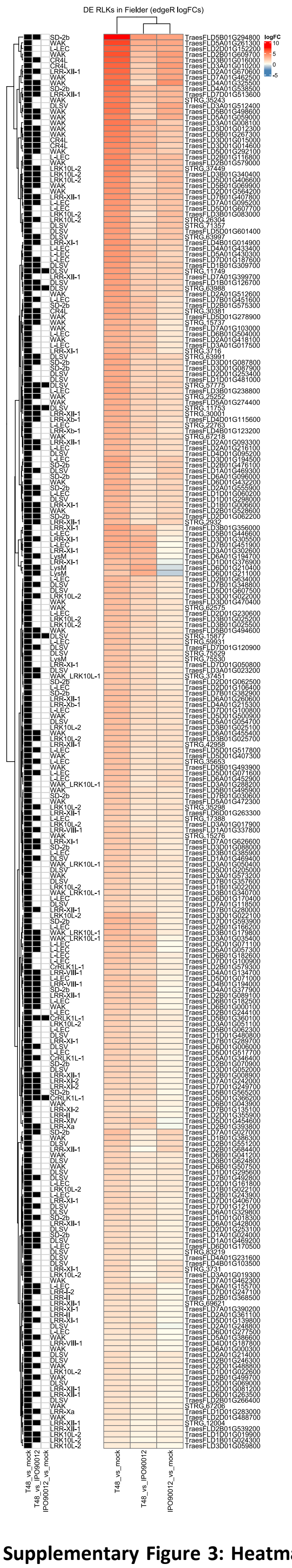

**Supplementary Figure 3: Heatmap of edgeR $log_2$-fold changes of all differentially expressed RLKs in the Fielder dataset.** RLKs were classified using iTAK (http://itak.feilab.net/cgi-bin/itak/online_itak.cgi). Filled squares indicate statistical significance in the respective contrast. Genes beginning with STRG are nTARs annotated using StringTie3.

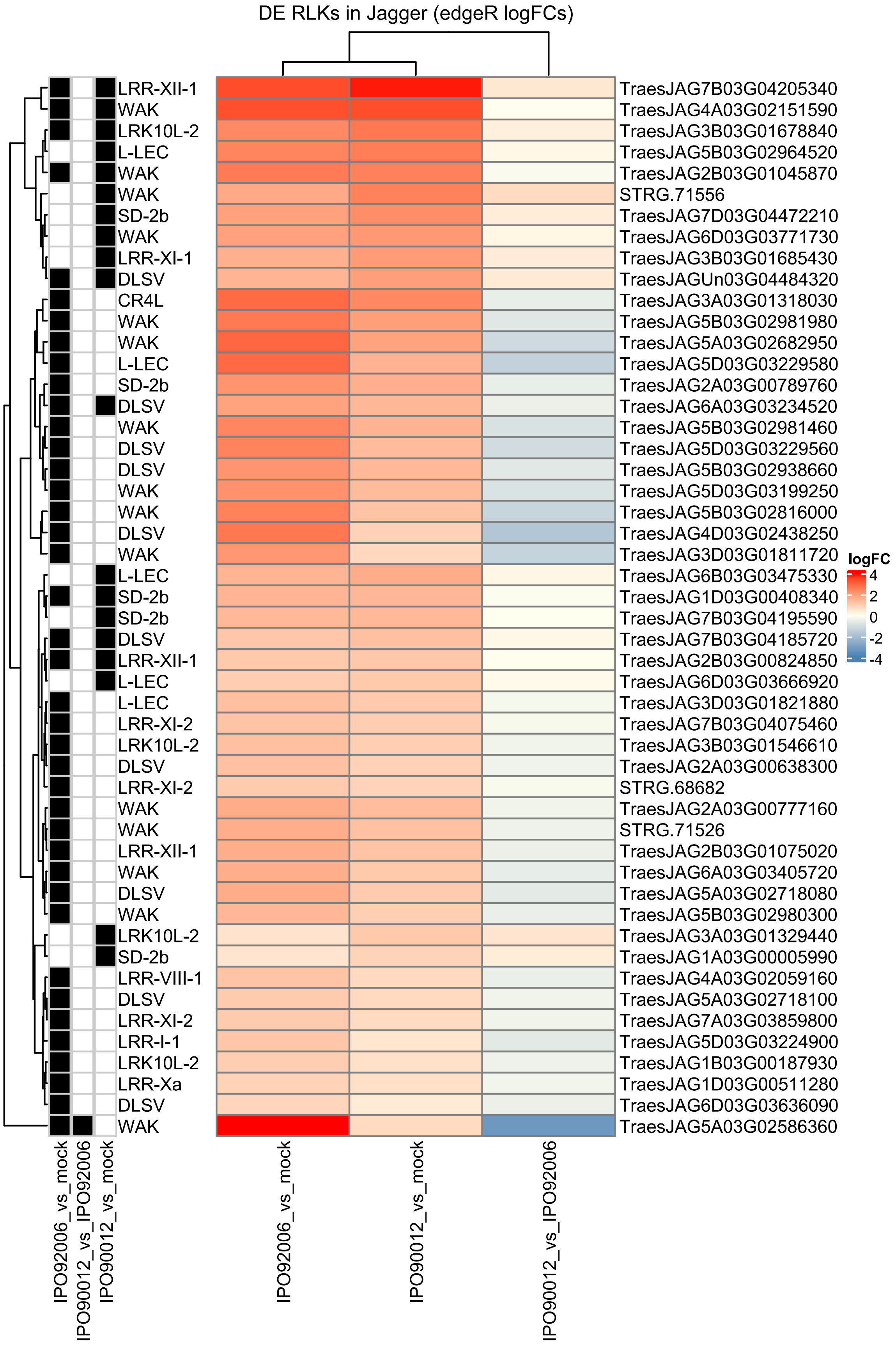


**Supplementary Figure 4: Heatmap of edgeR log2-fold changes of all differentially expressed RLKs in the Jagger dataset.** RLKs were classified using iTAK (http://itak.feilab.net/cgi-bin/itak/online_itak.cgi). Filled squares indicate statistical significance in the respective contrast. Genes beginning with STRG are nTARs annotated using StringTie3.

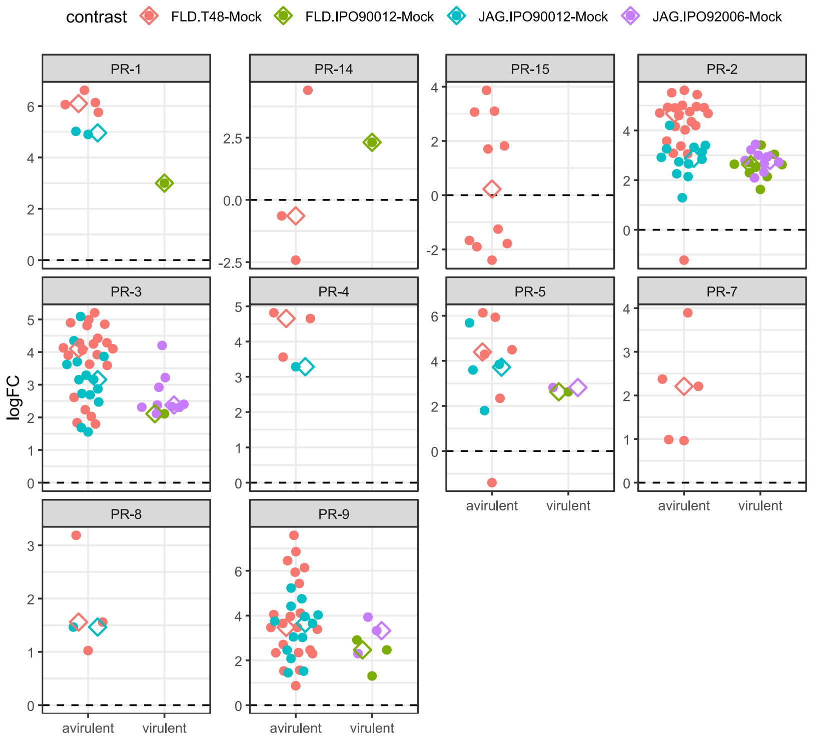


**Supplementary Figure 5: Number and expression changes (edgeR logFC) of wheat DEGs coding for different classes of pathogenesis-related proteins under infection with avirulent and virulent *Z. tritici* isolates.** Diamonds indicate mean logFC. PR proteins were classified by the detection of characteristic PFAM domains in InterProScan (Jones et al., 2014), and PR candidates were confirmed via BLASTp against a custom PR-protein database (*Supplementary Table S3*).

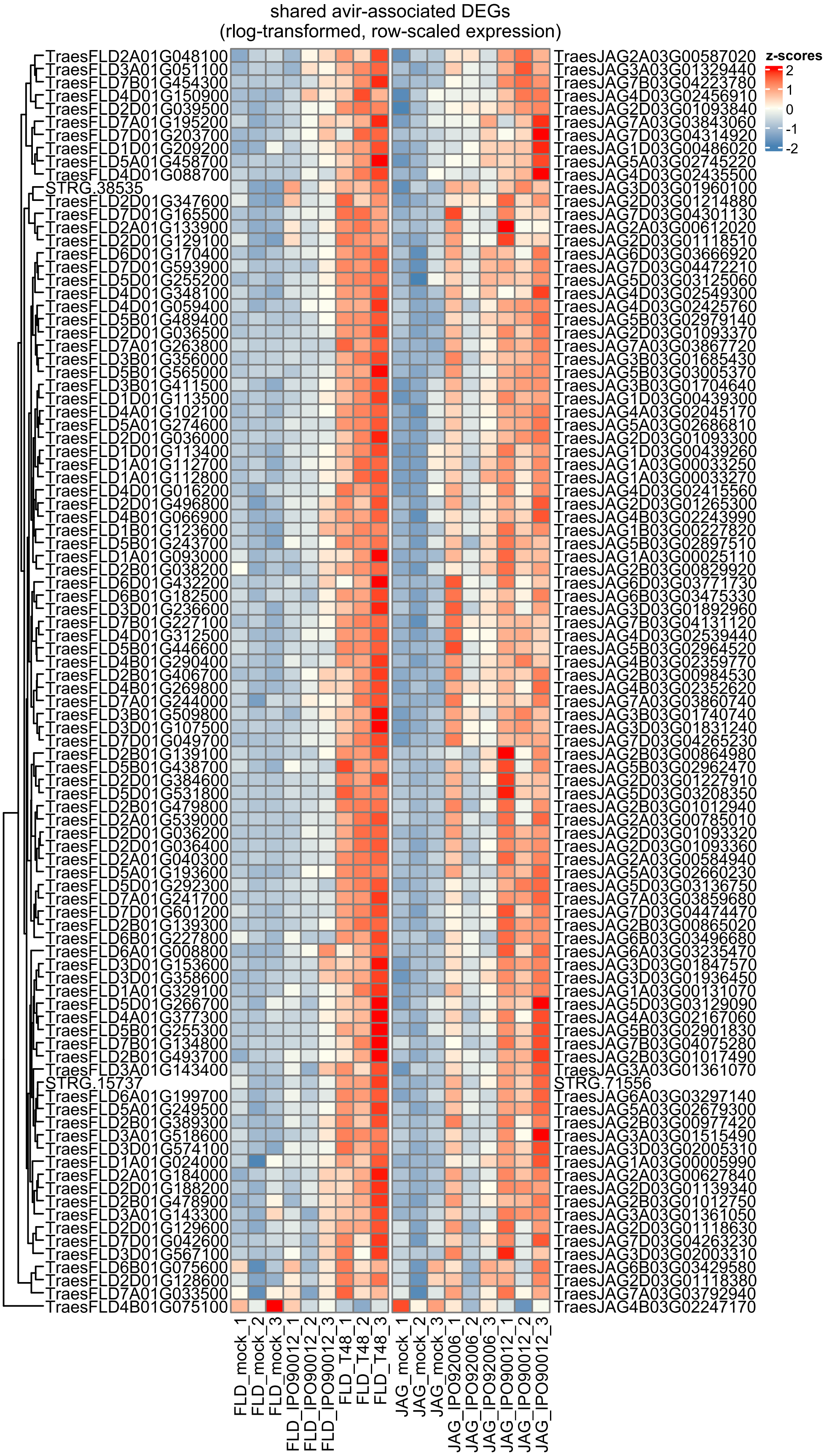


**Supplementary Figure 6: Expression of the 96 shared avirulence-associated DEGs (SAADs).** Expression values were regularized-log (rlog) transformed in DESeq2 and row-scaled for visualization. Numbers appended to sample names indicate biological replicates.

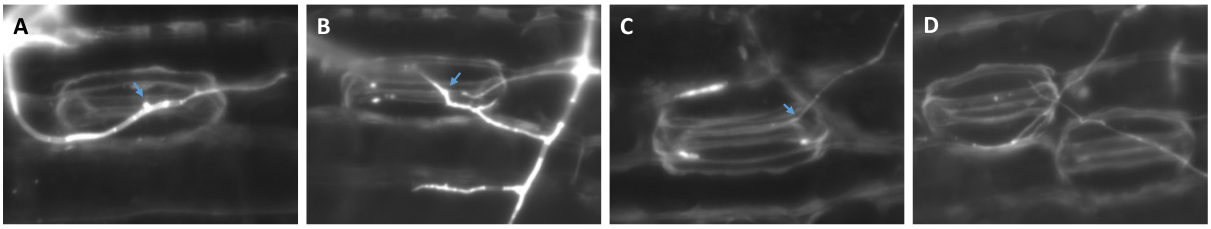


**Supplementary Figure 7: Representative microscopy images of *Z. tritici* hyphae interacting with wheat stomata.** (A-C) Stomata undergoing hyphal penetration attempts, highlighted with a blue arrow. (D) Hyphae contacting stomata without attempting penetration. (A) Observation of *Z. tritici* IPO92006 on Jagger at 6 dpi and (B-D) *Z. tritici* T48 on Fielder at 4 dpi.
