## Supplementary_Information_S1 for "Interaction-specific early wheat responses to *Zymoseptoria tritici* share enhanced defense-related transcription during incompatibility"

### Supplementary Information S1: Complementary analyses of the direct avirulent-isolate versus virulent-isolate (A–V) contrasts

#### Methods

For genome-wide comparison of the direct avirulent-isolate versus virulent-isolate responses, edgeR results were retained for all genes passing the original filterByExpr() filtering (see Experimental Procedures, "DEG testing with edgeR and DESeq2"), irrespective of statistical significance. Fielder and Jagger genes were matched using reciprocal best-hit CDS mappings, retaining 19,340 homologous gene pairs present in both filtered edgeR datasets. Genome-wide concordance was assessed descriptively using Pearson and Spearman correlations of edgeR $log_2FC$ estimates and by comparing effect direction between homologous pairs. To examine concordance among the strongest responses, genes were ranked independently within each cultivar using a signed quasi-likelihood statistic, calculated as (logFC) × √F, where F is the edgeR quasi-likelihood *F* statistic. For a series of percentile cut-offs, overlap between the corresponding positive or negative tails was calculated and expressed relative to the random expectation $k^2/N$ , where $k$ is the number of genes in each tail and $N$ is the total number of homologous genes. These analyses were used to assess genome-wide effect size and rank concordance rather than to define additional DEGs.

For the 96 SAADs, edgeR logFC estimates from the direct avirulent-isolate-versus-virulent-isolate contrasts were additionally extracted irrespective of FDR and compared descriptively between cultivars. These estimates were not used to define differential expression or membership of the SAAD set.

Genome-wide pre-ranked GO enrichment analyses were performed separately for Fielder and Jagger using the direct avirulent-isolate versus virulent-isolate edgeR contrasts. All genes retained in the corresponding filtered edgeR datasets and represented in the GO annotation universe (see Experimental Procedures, "Functional enrichment analyses of DEG sets") were included. Jagger genes were analyzed using Jagger gene identifiers, whereas Fielder genes were mapped to their Chinese Spring RefSeq v2.1 reciprocal best-hit homologues before matching to the Fielder GO annotation universe. Genes were ranked by the edgeR signed quasi-likelihood statistic (see above). GO terms were analysed separately for Biological Process, Molecular Function and Cellular Component using fgseaMultilevel from the fgsea Bioconductor package, with gene-set sizes restricted to 10–500 genes, gseaParam = 1, eps = 0 and nPermSimple = 10,000. Benjamini–Hochberg correction was applied within each ontology and cultivar. GO terms were considered concordantly enriched where they were significant at FDR ≤ 0.05 in both cultivars and showed enrichment in the same direction. As a sensitivity analysis, enrichment was repeated using logFC alone as the ranking statistic. Positive normalized enrichment scores (NES) indicate enrichment towards genes with higher estimated expression in the incompatible than in the compatible interaction, whereas negative NES values indicate enrichment in the opposite direction.

#### Results

##### 1. Genome-wide direct A–V expression effects show almost no overall gene-level concordance

Across 19,340 reciprocal best-hit Fielder–Jagger gene pairs passing the expression filter, direct A–V logFCs were essentially uncorrelated between cultivars (Pearson $r$ = 0.054; Spearman $\rho$ = −0.010), with only 49.5% of homologous pairs changing in the same direction. This provides strong evidence that there is no broad conserved gene-by-gene transcriptional program distinguishing the two incompatible from the corresponding compatible interactions (***Supplementary Figure 8***).

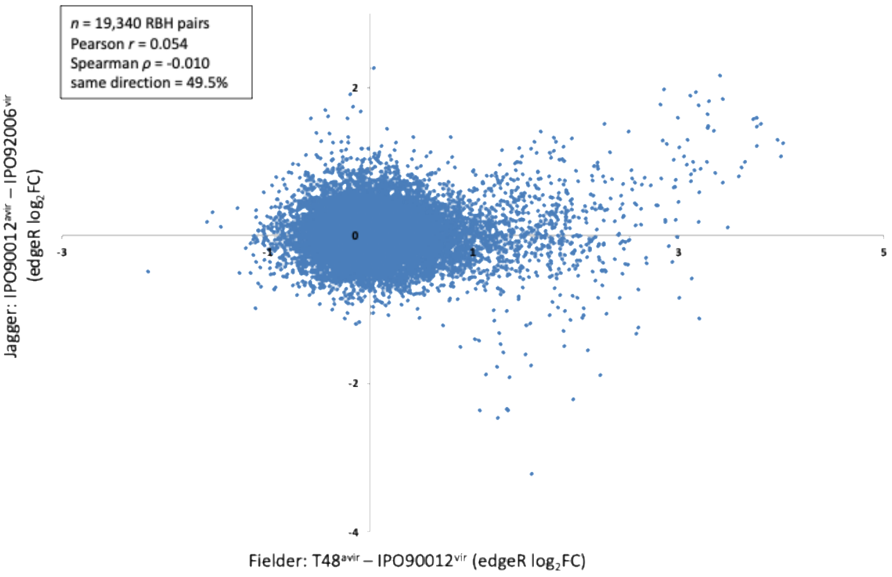


**Supplementary Figure 8. Genome-wide direct avirulent-isolate versus virulent-isolate expression effects show little overall concordance between Fielder and Jagger.** Each point represents one reciprocal best-hit Fielder–Jagger gene pair present in both filtered edgeR datasets ($n$ = 19,340). The x-axis shows the edgeR $\log_2$-fold change for Fielder plants infected with $T48^{avir}$ relative to those infected with $IPO90012^{vir}$, and the y-axis shows the corresponding $\log_2$-fold change for Jagger plants infected with $IPO90012^{avir}$ relative to those infected with $IPO92006^{vir}$. Positive values indicate higher estimated expression in the incompatible than in the compatible interaction. Vertical and horizontal reference lines indicate zero $\log_2$-fold change. Across all mapped gene pairs, overall concordance was weak (Pearson $r$ = 0.054; Spearman $\rho$ = −0.010), and 49.5% showed the same direction of effect in the two cultivars. Correlations are shown descriptively.

### 2. However, the strongest positive A–V responses are preferentially shared

When genes were ranked independently using sign(logFC) × $\sqrt{F}$, concordance became striking at the positive end of the distributions:

- top 1%: 38 shared homologous pairs versus 1.93 expected (19.7× enrichment)
- top 5%: 146 versus 48.35 expected (3.02×)
- top 10%: 297 versus 193.4 expected (1.54×)

Comparable enrichment was not observed among the strongest negative responses. Thus, convergence is concentrated specifically among genes showing the strongest increases in expression in the incompatible relative to the compatible interaction (***Supplementary Figure 9***).

The shared homologous gene pairs within the top 1%, 5% and 10% positive tails, together with ranks, effect estimates, $F$ statistics, FDRs, annotations and SAAD membership, are provided in ***Supplementary Table S5***.

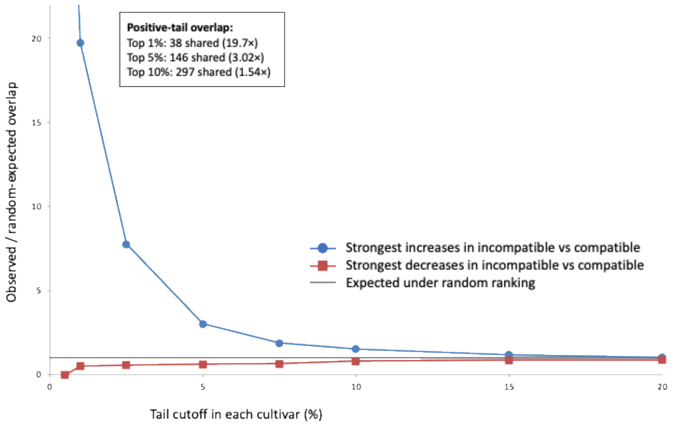


**Supplementary Figure 9. Genes showing the strongest increases in expression in incompatible relative to compatible interactions are preferentially shared between Fielder and Jagger.** The 19,340 reciprocal best-hit gene pairs present in both filtered edgeR datasets were ranked independently within each cultivar using a signed edgeR quasi-likelihood statistic, calculated as sign($\log_2$FC) × $\sqrt{F}$, where $F$ is the quasi-likelihood $F$ statistic. At each percentile cut-off, overlap between the corresponding tails of the Fielder and

Jagger rankings was calculated and expressed relative to the overlap expected under random ranking ($k^2/N$). Positive tails represent genes showing the strongest increases in expression in the incompatible relative to the compatible interaction, whereas negative tails represent genes showing the strongest decreases in expression in the incompatible relative to the compatible interaction. An observed/expected ratio of 1 indicates the overlap expected under random ranking. Overlap enrichments are presented descriptively because co-expression and gene-family structure mean that individual genes cannot be regarded as fully independent observations.

### 3. The independently defined SAAD set shows the same quantitative tendency

We also examined the direct A–V effect estimates for the 96 shared avirulence-associated DEGs (SAADs), which had originally been defined independently from the avirulent-isolate versus mock and virulent-isolate versus mock contrasts. 95 of the 96 SAADs had positive A–V effect estimates in both cultivars, i.e., higher estimated expression in the incompatible than in the compatible interaction. Their direct effects were also positively associated across cultivars (Pearson $R$ = 0.58). This comparison is deliberately descriptive because the SAADs were not defined using the direct A–V contrast, but it independently supports the idea of quantitative rather than qualitative convergence (***Supplementary Figure 10***).

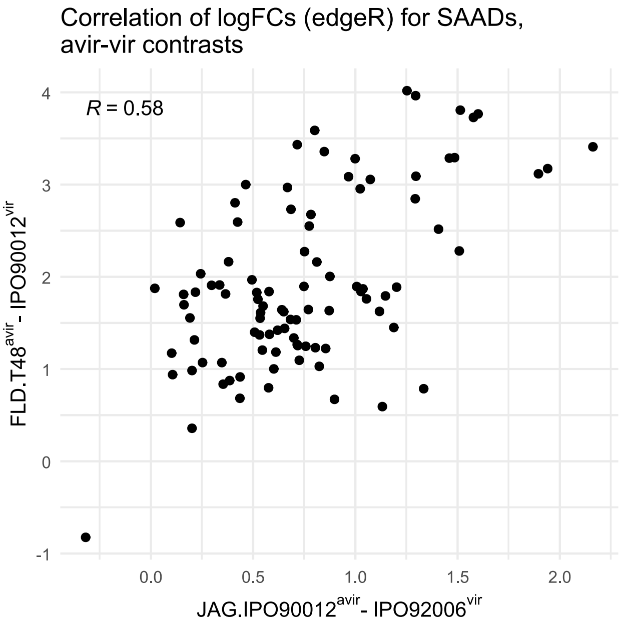


**Supplementary Figure 10. Direct avirulent-isolate versus virulent-isolate expression effect estimates for the 96 shared avirulence-associated DEGs (SAADs).** Each point represents one homologous SAAD pair. The x-axis shows the edgeR $\log_2$-fold change for Fielder plants infected with T48$^{avir}$ relative to those infected with IPO90012$^{vir}$, and the y-axis shows the corresponding $\log_2$-fold change for Jagger plants infected with IPO90012$^{avir}$ relative to those infected with IPO92006$^{vir}$. Positive values indicate higher estimated expression in the incompatible than in the compatible interaction. Ninety-five of the 96 SAADs showed positive effect estimates in both cultivars, while one showed negative effect estimates in both. Direct contrast effect estimates were positively associated between cultivars (Pearson $R$ = 0.58). Values are shown irrespective of FDR; the plot therefore visualizes the direction and magnitude of estimated effects and does not imply statistically significant differential expression between the two infected treatments for individual genes. Correlations are shown descriptively.

### 4. Genome-wide ranked GO analysis shows functional convergence

Finally, we performed independent fgseaMultilevel analyses of the two direct A–V contrasts. Despite the weak genome-wide gene-level correlation, we identified 22 GO terms significantly enriched in the same direction in both cultivars:

- 9 Biological Process (BP) terms
- 10 Molecular Function (MF) terms
- 3 Cellular Component (CC) terms

The concordantly enriched GO terms, together with cultivar-specific gene-set sizes, NES and FDR values and support from the $\log_2$FC-only sensitivity analysis, are summarized in ***Supplementary Table S6***. All nine concordant BP terms were positively enriched. The strongest common themes were chitin and cell-wall macromolecule catabolism, oxidative-stress and hydrogen-peroxide responses, cellular oxidant detoxification, glutathione metabolism, antifungal defense and carbohydrate metabolism. The MF results reinforce these processes, including chitinase activity, chitin binding, β-1,3-glucanase activity, peroxidase activity and glutathione transferase activity. The CC analysis additionally identified positive enrichment of the extracellular region, consistent with secreted/apoplastic defense-associated proteins. Most of the strongest functional signals were also recovered when $\log_2$FC alone was used for ranking, providing an independent sensitivity check (***Supplementary Figure 11***).

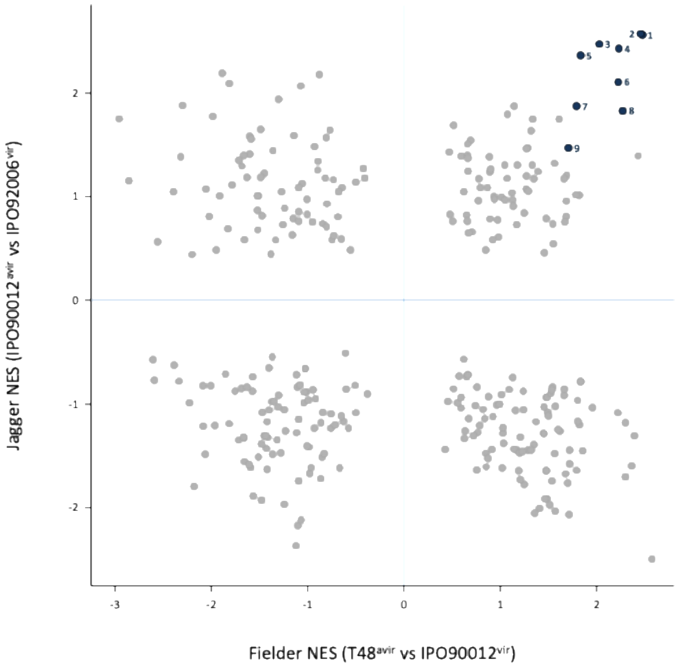


**Supplementary Figure 11. Concordant genome-wide ranked GO enrichment identifies shared defense-associated biological processes in incompatible interactions.** Each point represents one Biological Process GO term tested in both Fielder and Jagger ($n$ = 329). Genes were ranked independently within each cultivar using the signed edgeR test statistic, calculated as $\text{sign}(\log_2\text{FC}) \times \sqrt{F}$, from the direct avirulent-isolate versus virulent-isolate contrasts. The x-axis shows the normalized enrichment score (NES) for Fielder plants infected with T48$^{avir}$ relative to those infected with IPO90012$^{vir}$, and the y-axis shows the corresponding NES for Jagger plants infected with IPO90012$^{avir}$ relative to those infected with IPO92006$^{vir}$. Positive NES values indicate enrichment towards genes showing higher estimated expression in the incompatible than in the compatible interaction; negative values indicate enrichment in the opposite direction. Grey points show all common Biological Process terms. Deep-blue points indicate terms significantly enriched in the same direction in both cultivars at FDR ≤ 0.05. Numbered terms are: 1, chitin catabolic process; 2, cell wall macromolecule catabolic process; 3, response to oxidative stress; 4, hydrogen peroxide catabolic process; 5, cellular oxidant detoxification; 6, glutathione metabolic process; 7, toxin catabolic process; 8, defense response to fungus; and 9, carbohydrate metabolic process. All nine concordant significant terms showed positive enrichment in both cultivars.

Taken together, these four analyses substantially refine the story. The two incompatible interactions show very little genome-wide concordance at the level of individual homologous genes, arguing against a single conserved transcriptional program. Nevertheless, convergence becomes apparent in three complementary ways: among the strongest positive A–V gene responses, within the independently defined SAAD candidate set, and most clearly at the level of shared defense-associated biological functions. In other words, different cultivar–isolate combinations appear to recruit substantially different individual genes, but converge quantitatively on a restricted set of defensive outputs, particularly antifungal hydrolytic activity and redox/ROS homeostasis.
